# A Distal Super-Enhancer Regulates CD47 Expression and Promotes Macrophage-Mediated Phagocytosis of Ovarian Cancer Cells

**DOI:** 10.64898/2026.09.07.749763

**Authors:** Alok K. Mishra, Tanay Biswas, Anjan Roy, Shahid Banday, Arpit Katiyar, Anshul Sharma, Rui Li, Lihua Julie Zhu, Sunil K. Malonia, Bushra Ateeq

**Affiliations:** Department of Biological Sciences and Bioengineering, Indian Institute of Technology Kanpur, Kanpur, UP 208016, India; Department of Molecular Cell and Cancer Biology, UMass Chan Medical School, Worcester, MA 01605 USA; Frontier Research Institute of Interdisciplinary Sciences, Islamic University of Science and Technology, Awantipora 192122, J&K, India; Department of Biochemical Engineering, Harcourt Butler Technical University, Kanpur, Uttar Pradesh 208002, India; Department of Medicine, Division of Hematology and Oncology, Medical College of Wisconsin, Milwaukee, Wisconsin, USA; Mehta Family Center for Engineering in Medicine, Indian Institute of Technology Kanpur, Kanpur, UP 208016, India; Centre of Excellence for Cancer - Gangwal School of Medical Sciences and Technology, Indian Institute of Technology Kanpur, Kanpur, UP 208016, India

**Keywords:** HGSOC, CD47, super-enhancer, macrophage-mediated phagocytosis, innate immune evasion, fallopian tube origin, enhancer regulation, ovarian cancer

## Abstract

CD47 is an innate immune checkpoint that can protect tumor cells from macrophage-mediated clearance, but the regulatory mechanisms underlying its expression in ovarian cancer (OC) remain incompletely understood. Here, we investigated the epigenetic regulation of CD47 expression in OC. Single-cell and spatial transcriptomic analyses indicated preferential CD47 expression in malignant epithelial cells. Integration of publicly available epigenomic datasets revealed a distal regulatory region with enhancer-associated features across ovarian cancer models and HGSOC patient samples. A genome-wide CRISPR–Cas9 screen further implicated PITX2 and EP300 as regulators of CD47 expression. Chromatin immunoprecipitation assay showed PITX2 and EP300 occupancy at candidate enhancer elements within this CD47 regulatory region. Comparative epigenomic analyses suggested that features of this regulatory landscape are also present in fallopian tube epithelial cells and become more prominent in OC. suTogether, these findings support a PITX2-EP300 regulatory axis associated with CD47 expression and provide evidence for its potential contribution to tumor immune evasion.

**Highlights:**

1. A distal regulatory element is associated with CD47 expression in ovarian cancer.
2. PITX2 and EP300 regulate CD47 expression through a distal enhancer.
3. PITX2 or EP300 depletion enhances macrophage-mediated phagocytosis.
4. CD47 enhancer activity pre-exists in fallopian tube epigenome.

## Introduction

Tumor immune evasion is a fundamental hallmark of cancer, allowing malignant cells to evade both innate and adaptive immune surveillance and thereby sustain tumor progression^1–3^. CD47 functions as a major innate immune checkpoint by engaging SIRPα on macrophages to inhibit phagocytic clearance^4–6^. Although CD47 is widely expressed in normal tissues and contributes to recognition of self, many cancers exploit this pathway by upregulating CD47 to evade macrophage-mediated clearance ^5–7^. Elevated CD47 expression has been reported across multiple malignancies, including ovarian cancer, where increased expression has been associated with aggressive disease and adverse clinical outcomes^4,5,7–12^. The transcriptional regulation of CD47 transcription is regulated by multiple context-dependent signaling pathways and transcriptional programs. Studies of CD47 transcription have largely focused on promoter-proximal regulatory mechanisms, with transcription factors including α-Pal/NRF-1^13^, MYC ^14^, HIF-1α^15^, and NF-κB^16^, shown to regulate CD47 transcription in response to metabolic, oncogenic, inflammatory, and hypoxic signals. ^6,14–18^. These studies have established promoter-proximal regulation as an important mechanism controlling CD47 expression. More recently, Betancur et al. ^19^ demonstrated that CD47 expression can also be regulated by distal enhancer elements, including a super-enhancer. Their study identified a downstream super-enhancer associated with CD47 transcription in breast cancer, as well as distinct upstream enhancer or super-enhancer elements in T-cell acute lymphoblastic leukemia (T-ALL) and diffuse large B-cell lymphoma (DLBCL), highlighting tumor-type-specific distal regulation of CD47 ^19^. Whether such enhancer-dependent regulation operates in other solid tumors and how these regulatory elements are established during tumorigenesis remain important unanswered questions. HGSOC provides a compelling context in which to address these questions because its development is closely linked to lineage-specific transcriptional and epigenomic programs^20–27^. Recent epigenomic studies have established that HGSOC exhibits widespread enhancer remodeling and lineage-specific transcriptional circuitry ^24,25^. Given that CD47 is frequently overexpressed in HGSC and functions as a major innate immune checkpoint, defining the cis-regulatory mechanisms governing its expression may reveal previously unrecognized regulatory dependencies that could be exploited to modulate tumor immune recognition.

Here, we integrate publicly available transcriptomic and epigenomic datasets with genome-wide CRISPR-based functional genomics to identify regulatory elements and transcriptional regulators associated with CD47 expression in ovarian cancer. We sought to define the enhancer landscape underlying aberrant CD47 activation, determine whether it arises de novo during tumorigenesis or is co-opted from the normal cell of origin and identify the molecular regulators. Together, our findings uncover an epigenetic mechanism regulating CD47 expression and driving innate immune checkpoint activation in ovarian cancer.

## Results

### CD47 expression is associated with a distal enhancer landscape in ovarian cancer

Although CD47 is broadly expressed across normal tissues, its dysregulated expression has been reported in multiple malignancies including ovarian cancer ^7,8,17,18^. Consistent with previous observations, comparative analysis of TCGA and GTEx transcriptomic datasets identified ovarian cancer as displaying among the highest aberrant CD47 overexpression relative to normal ovarian tissue across TCGA malignancies (**Fig. S1A**). To determine the cellular distribution of CD47 expression in high-grade serous ovarian cancer (HGSOC), we interrogated seven independent single-cell RNA-sequencing datasets from the TISCH2 database, encompassing diverse patient cohorts and tumor microenvironments.^28^ Across these independent cohorts, CD47 expression was consistently enriched in malignant epithelial cells compared with stromal and immune cell populations. (**Fig. 1A-B and S1B-C left)**. As expected, the cognate receptor *SIRPA* was predominantly expressed in monocyte/macrophage populations (**Fig. S1B-C, right)**. Independent spatial transcriptomic analysis of the Broad Institute HGSOC spatial cohort (SCP2640) further confirmed preferential localization of CD47 expression within malignant tumor regions, while stromal and immune compartments displayed comparatively lower expression, supporting tumor cell-intrinsic expression of this innate immune checkpoint (**Fig. S1D-E**).

**Figure 1.**
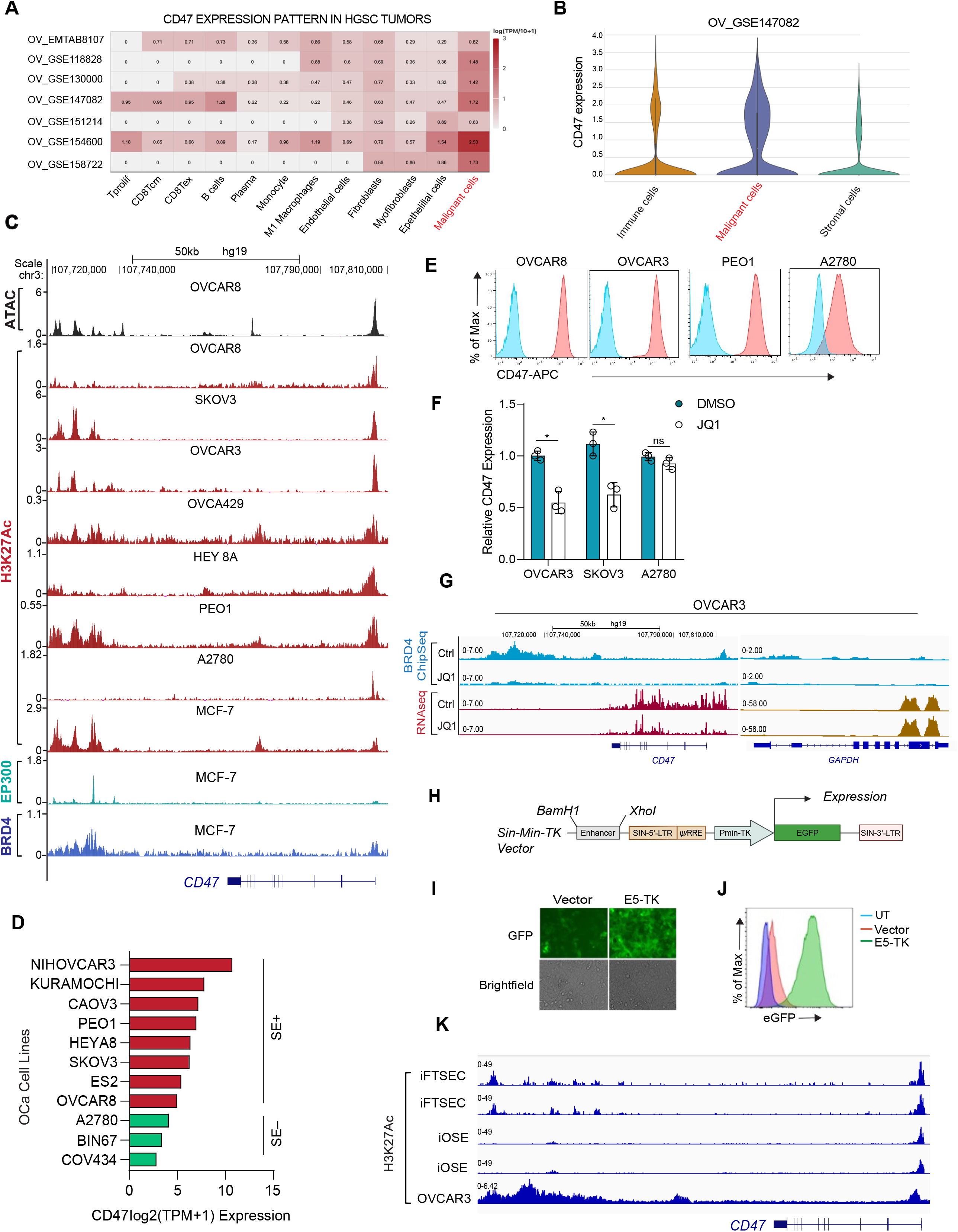
CD47 is preferentially expressed in malignant epithelial cells and is associated with an active enhancer landscape in HGSOC. (A) Heatmap showing CD47 expression across major cell populations in multiple publicly available single-cell RNA-seq datasets of ovarian cancer, demonstrating preferential CD47 expression in malignant cells compared with non-malignant stromal and immune populations. (B) Violin plot showing CD47 expression across immune, malignant, and stromal cell populations in the OV_GSE147082 dataset. (C) Genome-browser tracks displaying ATAC-seq and H3K27ac signals from ovarian cancer cell lines and p300/BRD4 occupancy from MCF7 cells across the CD47 locus (D) CD47 transcript expression across a panel of ovarian cancer cell lines, with cell lines classified according to the presence (SE+) or absence (SE−) of a CD47-associated super-enhancer. (E) Flow-cytometry histograms showing surface CD47 expression in OVCAR8, OVCAR3, PEO1, and A2780 ovarian cancer cells. (F) Bar graph showing qRT-PCR analysis of CD47 transcription upon JQ1 treatment (G) Genome-browser tracks showing BRD4 occupancy at the CD47 locus following treatment with the BET inhibitor JQ1 or DMSO control, together with RNAseq track showing CD47 expression in OVCAR3 following JQ1 treatment. (H) Schematic representation of the Sin-minTK enhancer reporter vector. (I) Fluorescence micrographs showing GFP expression from the E5-PminTK enhancer reporter. (J) Flow cytometry histograms showing GFP expression from the E5-PminTK enhancer reporter. (K) CD47 enhancer landscape across ovarian cancer precursor and HGSOC cells; IGV tracks showing H3K27ac occupancy across the CD47 genomic region in fallopian tube secretory epithelial cells (iFTSEC), immortalized ovarian surface epithelial cells (iOSE), and OVCAR3 cells. The H3K27ac landscape demonstrates conservation of the CD47 super-enhancer region from the fallopian tube epithelial lineage to HGSOC.

We next asked whether the elevated CD47 expression in ovarian cancer is associated with a distal cis-regulatory element. A seminal study by Betancur et al. has identified CD47-associated super-enhancer located downstream of the CD47 gene in breast cancer and demonstrated that this region contains several constituent enhancers, including the most functionally active enhancer in MCF7 ^19^. Because enhancer usage can be context-and cancer-type-dependent, we used this previously characterized CD47 regulatory region as a genomic reference to investigate CD47 associated superenhancer in OC. Integration of publicly available epigenomic datasets identified a conserved regulatory region located approximately 100 kb downstream of the CD47 transcription start site (TSS) in ovarian cancer models. The candidate region exhibited increased chromatin accessibility, with a prominent ATAC-seq peak in OVCAR8 cells and was associated with robust H3K27ac enrichment across multiple ovarian cancer cell lines, including OVCAR8, OVCAR3, OVCA429, HEY8A, PEO1, and SKOV3. However, interestingly, A2780, BIN67 and COV434 cells displayed little or no H3K27ac enrichment at the corresponding locus **(Fig. 1C and S2A**). These findings indicate that this distal regulatory region exhibits an active chromatin state in a subset, but not all, ovarian cancer cell lines. MCF7 p300 and BRD4 occupancy at the orthologous region provided a reference for the previously characterized CD47 super-enhancer (**Fig. 1C**). The coincident enrichment of H3K27ac, chromatin accessibility, EP300, and BRD4 is consistent with the molecular features of an active enhancer-rich regulatory domain.

We next examined whether the presence of this regulatory region was associated with CD47 expression. Integration of DepMap transcriptomic data with available H3K27ac ChIP-seq profiles revealed that ovarian cancer cell lines harboring the CD47-associated regulatory region exhibited higher CD47 transcript levels than cell lines lacking detectable enhancer marks at this locus (**Fig. 1D, Table S1**). We then assessed CD47 at the protein level by flow cytometry. Consistent with the transcriptomic and epigenomic analyses, ovarian cancer cell lines with an active regulatory region, including OVCAR8, SKOV3, and PEO1, displayed substantially higher cell-surface CD47 expression than the superenhancer-negative A2780 cell line (**Fig. 1E**). These observations support an association between enhancer-associated activity at the distal CD47 regulatory region and CD47 expression in ovarian cancer cells.

Because BRD4 is a major transcriptional coactivator associated with active enhancer and super-enhancer domains, we next asked whether CD47 expression depended on BRD4 activity. Pharmacological inhibition of BET bromodomains with JQ1 treatment significantly reduced CD47 transcript levels harboring the active CD47 regulatory region, whereas the effect was substantially less pronounced in the enhancer-negative A2780 cells (**Fig. 1F**). To determine whether this response was accompanied by changes in BRD4 occupancy at the CD47 regulatory region, we analyzed publicly available BRD4 ChIP-seq data from OVCAR3 cells before and after JQ1 treatment. JQ1 treatment resulted in a marked reduction of BRD4 occupancy at the CD47 regulatory region which was accompanied by a reduction in CD47 RNA-seq signal (**Fig. 1G and S2B**). These findings support a role for BRD4-associated enhancer activity in maintaining CD47 expression, particularly in super-enhancer positive ovarian cancer cells.

Next, we sought to directly test whether the conserved regulatory sequence identified at the CD47 locus possesses intrinsic enhancer activity in ovarian cancer cells. Because the previously characterized E5 constituent enhancer resides within the downstream CD47 regulatory region defined in MCF7 cells, we used the same E5 sequence and primer set described by Betancur et al. ^19^, to experimentally assess its activity in ovarian cancer. The E5 region was cloned upstream of a minimal *thymidine kinase* promoter in an enhancer-reporter construct and expressed in OVCAR8 cells. The E5-containing reporter showed increased activity relative to the minimal-promoter control, demonstrating intrinsic enhancer activity of this conserved sequence in OVCAR8 cells **(Fig. 1H-J**).

Given that HGSOC is thought to arise from fallopian tube secretory epithelial cells (FTSECs)^29^, we asked whether SE-associated chromatin features at the CD47 locus are detectable in nonmalignant fallopian tube epithelial cells or aquired *de novo* during OC evolution (**Fig. S3**). Comparison of H3K27ac ChIP-seq profiles revealed a reproducible distal H3K27ac-marked enhancer at the CD47 locus in two iFTSEC biological replicates, with a weaker corresponding signal in iOassSE cells. In contrast, OVCAR3 cells displayed broader and stronger H3K27ac enrichment across the same region, consistent with cancer-associated enhancer remodeling and expansion (**Fig 1K**). These findings suggest that HGSOC may co-opt and expand a pre-existing lineage-associated regulatory element at the CD47 locus, rather than establishing the enhancer *de novo* during transformation.

### Genome-wide CRISPR screen identifies candidate regulators of CD47 expression in ovarian cancer

To systematically identify factors regulating CD47 expression, we performed a genome-wide CRISPR-Cas9 loss-of-function screen in the ovarian cancer cell line OVCAR8, using cell-surface CD47 abundance as the phenotypic readout (**Fig. 2A**). OVCAR8 cells expressing Cas9 were transduced with the Brunello genome-wide sgRNA library, comprising approximately 76,000 sgRNAs targeting ∼19,000 protein-coding genes, to enable systematic loss-of-function interrogation of genes involved in CD47 surface expression. Following library transduction and selection, cells were stained with the anti-CD47 monoclonal antibody (clone;CC2C6), which recognizes the N-terminal pyroglutamylated form of CD47, and subjected to fluorescence-activated cell sorting to isolate cells with reduced cell-surface CD47 abundance. In parallel, an unsorted cell population was collected to provide a reference for sgRNA representation in the input population. Genomic DNA was isolated from the sorted and unsorted populations, sgRNA sequences were amplified and quantified by deep sequencing, and candidate genes were identified based on their differential representation in the CD47-low population relative to the unsorted control (**Table S2**). CD47 itself emerged as the top-ranked hit, providing an internal positive control for the screen and supporting the robustness of the surface-CD47-based selection strategy. Notably, QPCTL, the enzyme responsible for catalyzing the N-terminal pyroglutamylation of CD47 ^30^ was also among the top-ranked candidates. Because recognition of CD47 by the CC2C6 antibody depends on this post-translational modification, the identification of QPCTL provided an orthogonal internal control that further supported the specificity of the screening phenotype. Together, these findings indicated that the screen effectively captured both CD47 itself and factors required for its antibody-detectable cell-surface state.

**Figure 2.**
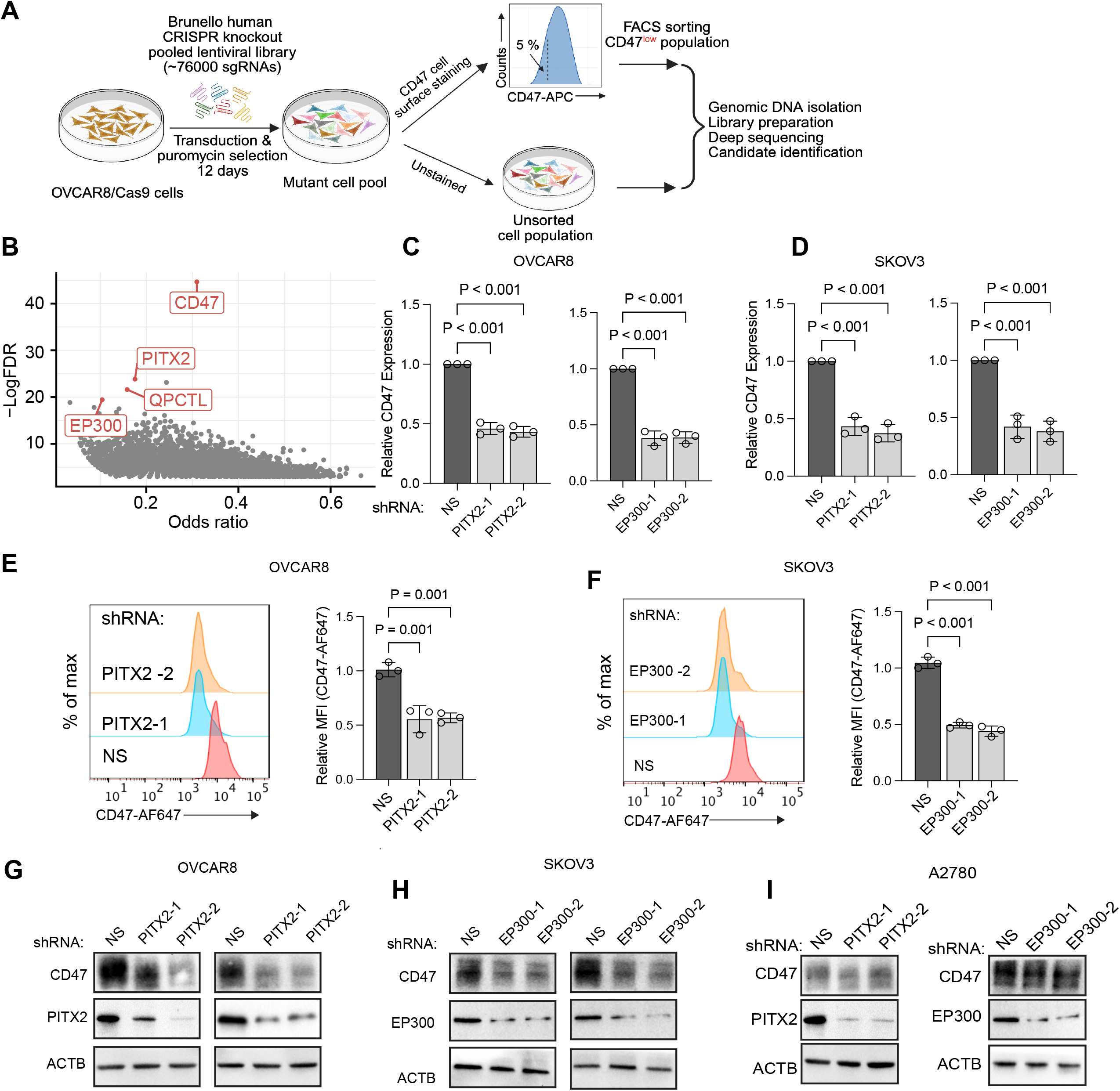
Genome-wide CRISPR screen identifies PITX2 and EP300 as candidate regulators of CD47 expression. (A) Schematic of the genome-wide CRISPR/Cas9 knockout screen performed in Cas9-expressing OVCAR8 cells. (B) Volcano plot showing candidate genes identified from the cell-surface CD47 screen. CD47, QPCTL, PITX2, and EP300 were among the top-ranked hits. (C-D) Relative CD47 mRNA expression following independent shRNA-mediated depletion of PITX2 or EP300 in OVCAR8 (C) and SKOV3 (D) cells. Two independent shRNAs targeting each gene were used, with a non-silencing (NS) shRNA as the control. (E-F) Representative flow cytometry histograms and quantification of cell-surface CD47 following PITX2 (E) or EP300 (F) knockdown in OVCAR8 cells. Cell-surface CD47 abundance was quantified as mean fluorescence intensity (MFI). (G-I) Immunoblot analysis of CD47 protein expression following PITX2 or EP300 knockdown in OVCAR8 (G), SKOV3 (H), and A2780 (I) cells. ACTB was used as a loading control. Quantitative data are presented as mean ± SEM where applicable; statistical significance is indicated in the individual panels.

Among the candidate regulators, the paired-like homeodomain transcription factor PITX2 and the histone acetyltransferase EP300 emerged as significant hits (**Fig. 2B, Table S3**). To independently validate these candidates, we performed shRNA-mediated knockdown of PITX2 or EP300 using two independent shRNAs in multiple ovarian cancer cell lines and assessed CD47 expression by qRT-PCR, flow cytometry, and immunoblotting. Knockdown of either PITX2 or EP300 consistently reduced CD47 mRNA and total protein levels and was accompanied by decreased cell-surface CD47 expression (**Fig 2C-H**). Notably, CD47 expression was not significantly altered in A2780 cells, suggesting that PITX2-EP300 regulation of CD47 is context dependent (**Fig 2I**).

### PITX2 and EP300 occupy a distal enhancer associated with CD47 expression

Given the established roles of PITX2 and EP300 in enhancer regulation and chromatin accessibility across diverse cellular contexts ^31,32^, we subsequently investigated whether PITX2 and EP300 are associated with the distal CD47 regulatory region. First we performed a co-immunoprecipitation assays in cells overexpressing FLAG-tagged PITX2 and HA-tagged EP300. Reciprocal immunoprecipitation detected a physical interaction between PITX2 and EP300 under the experimental conditions (**Fig. 3A**). Inspection of the distal CD47 regulatory region identified three putative PITX2-binding motifs (SE1-SE3) within regions exhibiting strong H3K27ac enrichment and chromatin accessibility. Motif analysis identified consensus PITX2 recognition sequences within these candidate enhancer elements, with SE2 and SE3 overlapping prominent H3K27ac and ATAC-seq peaks (**Fig. 3B**). To determine whether PITX2 occupies the CD47 enhancer in ovarian cancer cells, we performed chromatin immunoprecipitation followed by quantitative PCR (ChIP-qPCR) across the three enhancer regions and the CD47 promoter. PITX2 occupancy was detected at all three candidate enhancer regions (**Fig. 3C**). EP300 was also enriched at these candidate enhancer elements, whereas minimal enrichment was detected at the promoter region (**Fig. 3D**). Moreover, H3K27ac enrichment was higher across the candidate enhancer regions than at the promoter, consistent with an active enhancer-associated chromatin state (**Fig. 3E**). Collectively, these findings demonstrate that PITX2 and EP300 are enriched at the same distal enhancer region downstream of CD47 that is characterized by enhancer-associated histone acetylation and accessible chromatin architecture.

**Figure 3.**
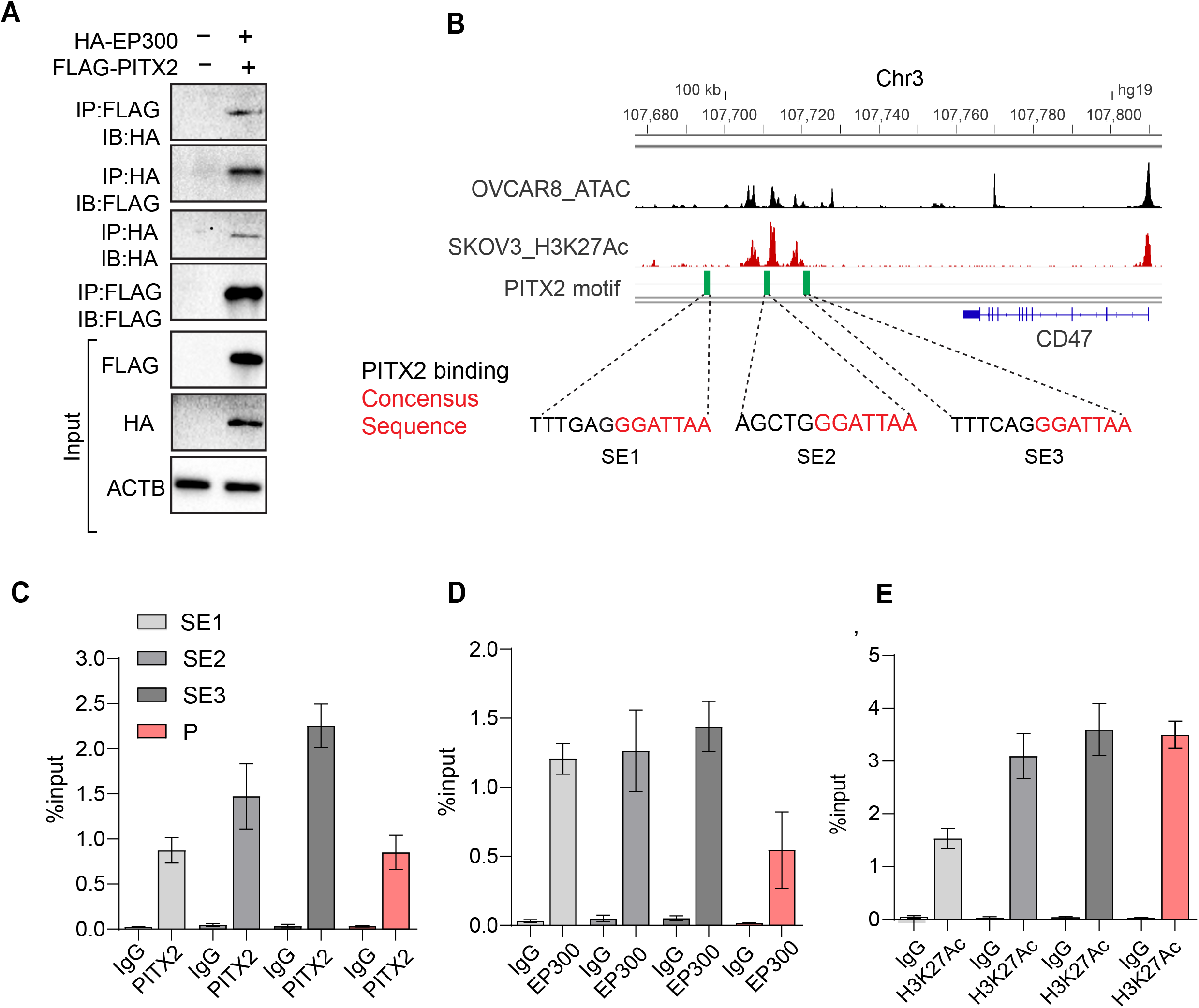
PITX2 interacts with EP300 and occupies an active enhancer region at the CD47 locus. (A) Co-immunoprecipitation demonstrating interaction between PITX2 and EP300 using FLAG-PITX2 and HA-EP300 overexpression. Reciprocal immunoprecipitation was performed using anti-FLAG or anti-HA antibodies and immunoblotting for the indicated proteins. (B) Genome-browser view of the CD47 locus on chromosome 3 (hg19), showing OVCAR8 ATAC-seq and SKOV3 H3K27ac signals across a distal ∼100-kb regulatory region downstream of CD47. Three candidate enhancer elements (SE1-SE3) contain consensus PITX2-binding motifs. (C-E) ChIP-qPCR using antibodies against endogenous PITX2 (C), EP300 (D), and H3K27ac (E) showing enrichment at the indicated CD47 enhancer regions (SE1, SE2, and SE3) and promoter (P).

### PITX2 and EP300 regulate CD47 expression and macrophage-mediated clearance in ovarian cancer

To determine whether PITX2-or EP300-dependent regulation of CD47 influences macrophage-mediated clearance, we examined the functional consequences of depleting either factor. Genetic depletion of PITX2 or EP300 significantly increased macrophage-mediated engulfment of ovarian cancer cells compared with control cells and was accompanied by reduced CD47 expression **(Fig. 4A-B).** These findings support a functional association between PITX2/EP300-dependent CD47 expression and susceptibility of ovarian cancer cells to macrophage-mediated clearance.

**Figure 4.**
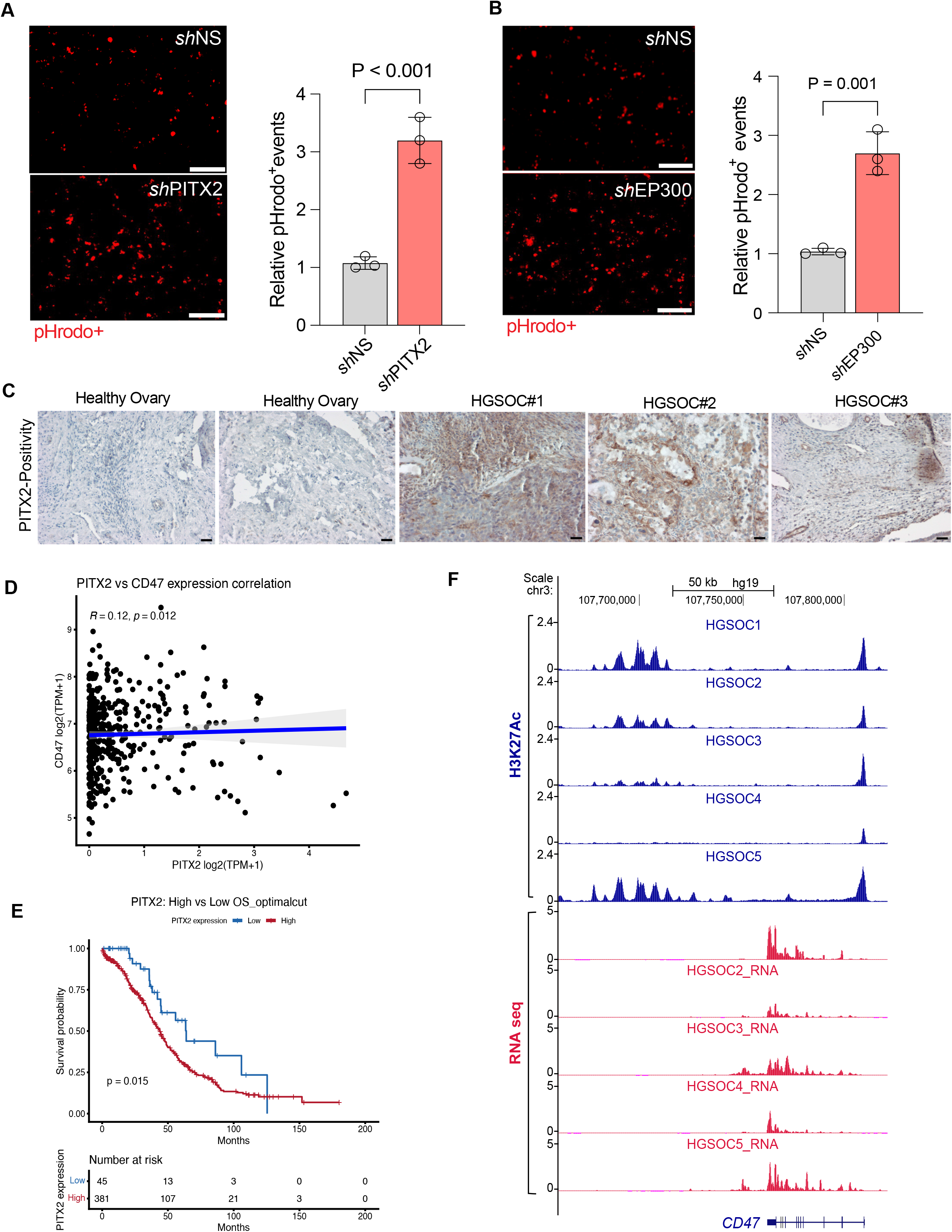
PITX2 or EP300 depletion enhances macrophage phagocytosis, and enhancer-driven CD47 expression is clinically relevant in HGSOC (A) Representative images and quantification of macrophage-mediated phagocytosis assays using pHrodo-labeled ovarian cancer cells expressing control shRNA (shNS) or shPITX2. Phagocytic events were quantified as relative pHrodo-positive events. Data are presented as mean ± SD from independent experiments. Statistical significance was determined using a Student’s *t*-test. (B) Representative images and quantification of macrophage-mediated phagocytosis assays following EP300 depletion (shEP300) compared with shNS controls. Relative pHrodo-positive events are shown as mean ± SD. Statistical significance was assessed using Student’s *t*-test (C) Representative immunohistochemical staining for PITX2 in normal ovarian tissue and primary high-grade serous ovarian cancer (HGSC) specimens. Increased nuclear PITX2 expression was observed in HGSC samples relative to healthy ovarian tissue. (D) Correlation analysis between PITX2 and CD47 transcript abundance in TCGA ovarian cancer samples. (E) Kaplan-Meier overall survival analysis of HGSOC patients stratified by PITX2 expression. High PITX2 expression was associated with shorter overall survival. (F) H3K27ac enrichment at the distal CD47 regulatory region in five independent HGSOC specimens and its association with CD47 transcript abundance using matched RNA-seq data.

We next examined the association between PITX2 expression and the clinical and molecular features of HGSOC. Immunohistochemical analysis of representative HGSOC specimens showed increased nuclear PITX2 staining compared with the normal ovarian tissues examined, consistent with the elevated PITX2 mRNA expression observed in TCGA datasets (**Fig. 4C, S4A**). Consistent with the experimental evidence linking PITX2 to CD47 expression, analysis of publicly available TCGA-HGSC data showed a modest positive correlation between *PITX2* and *CD47* transcript abundance (**Fig. 4D**). We next examined the association between *PITX2* expression (log2[TPM+1]) and overall survival in 426 HGSOC patients with matched clinical and RNA-seq data. *PITX2* expression was dichotomized into high and low groups using an optimal cutpoint. Patients with high *PITX2* expression showed significantly shorter overall survival (log-rank *p* = 0.015) (**Fig. 4E**).

We then asked whether the enhancer-associated chromatin features identified in ovarian cancer cell lines are retained in primary HGSOC tumors. Analysis of publicly available H3K27ac ChIP-seq datasets from five independent HGSOC specimens revealed H3K27ac enrichment at the distal CD47 regulatory region identified in ovarian cancer cell lines. Integration with matched RNA-seq data showed that greater H3K27ac enrichment at this locus was associated with higher CD47 transcript abundance (**Fig. 4F**). These observations support the presence of an active enhancer-associated chromatin state at the CD47 locus in primary HGSC tumors and its association with CD47 expression. Collectively, these findings link PITX2 and EP300 to CD47 expression and macrophage-mediated clearance and show that the enhancer-associated chromatin landscape identified in ovarian cancer cell lines is also observed in primary HGSOC tumors.

## Discussion

CD47 is widely expressed in normal tissues and has been considered a broadly expressed “don’t-eat-me” signal; however, accumulating evidence indicates that its expression is not transcriptionally uniform across tumors^6–8,10–12,33^. Rather, malignant cells can acquire or activate tumor-context-specific cis-regulatory elements that reinforce CD47 expression^19^. Here, we identify a distal CD47 regulatory region associated with CD47 expression in HGSOC and define PITX2 and EP300 as regulators of this expression program. We found that a region located approximately 100 kb downstream of the CD47 transcriptional start site displays multiple characteristics of an active enhancer/super-enhancer, including H3K27ac and H3K4me1 enrichment, increased chromatin accessibility and BRD4 occupancy. Reporter assay further demonstrated that the conserved E5 sequence possesses intrinsic enhancer activity in ovarian cancer cells, supporting its potential contribution to CD47 regulation. Thus, our findings place CD47 within a tumor-associated cis-regulatory circuit in HGSOC and provide a mechanistic insight for how elevated CD47 expression can be maintained at the chromatin level.

A particularly important aspect of our study is the identification of PITX2 as a regulator of CD47 expression and its occupancy at the distal CD47 regulatory region. PITX2 is a conserved paired-like homeobox transcription factor with well-established roles in developmental patterning and cell-fate specification. During embryogenesis, Nodal signaling induces PITX2 through a dedicated enhancer, after which PITX2 activates downstream transcriptional programs that control tissue specification and organ morphogenesis.^34–37^. PITX2 has previously been implicated in ovarian cancer biology, where its expression is increased in ovarian tumors and has been associated with tumor growth, migration, invasion, and activation of oncogenic signaling pathways^38,39^. However, the mechanisms through which PITX2 contributes to immune evasion have remained undefined.

Our findings provide a mechanistic link between PITX2 and an innate immune checkpoint by demonstrating PITX2 occupancy at the distal CD47 regulatory region together with enrichment of the transcriptional coactivator EP300. EP300/CBP are established components of active enhancer machinery and are frequently associated with lineage-and cell-state-specific enhancers, where their acetyltransferase activity contributes to H3K27ac deposition and enhancer activation^32^. Importantly, however, our data do not establish whether PITX2 directly recruits EP300 to the CD47 enhancer, whether both factors are independently recruited by other transcriptional regulators, or whether their occupancy reflects assembly of a larger enhancer-associated complex. This distinction is relevant because PITX2 has been shown in other biological contexts to associate with distinct chromatin regulators and transcriptional cofactors, including CBP and the MLL4/KMT2D complex, demonstrating that its transcriptional activity can be mediated through context-dependent interactions with chromatin-modifying machinery^40,41^. Thus, while our findings are suggestive of a model in which PITX2 contributes to establishment or maintenance of an EP300-associated active enhancer state at CD47, however, whether PITX2 directly recruits EP300 or other histone acetyltransferases remains to be determined.

Our findings are also consistent with the broader regulatory organization of HGSOC, in which lineage-associated transcription factors occupy extensive enhancer and super-enhancer landscapes^24–26^. Recent studies have identified transcription factors such as PAX8, SOX17, and MECOM as components of HGSOC core regulatory circuitry, with these factors occupying super-enhancer-associated regions and coordinating transcriptional programs that define malignant epithelial cell identity. Such observations support the concept that HGSOC is driven not only by recurrent genetic alterations but also by lineage-specific transcriptional and enhancer programs.^24^

Our epigenomic analysis further provides an important link between the CD47 enhancer and the emerging understanding of HGSOC cell of origin. HGSOC is now widely considered to arise predominantly from secretory epithelial cells of the distal fallopian tube, with p53 signatures and serous tubal intraepithelial carcinoma (STIC) representing recognized precursor lesions in a substantial fraction of cases. Genomic analyses have demonstrated evolutionary relationships between STIC lesions and concurrent HGSOC, while experimental lineage-tracing and transformation studies have demonstrated the capacity of fallopian-tube epithelial cells to give rise to HGSOC-like tumors. At the same time, experimental studies indicate that ovarian surface epithelium can also serve as a cell of origin, emphasizing that HGSOC likely represents a biologically heterogeneous disease with more than one possible cellular origin^20,21,27,42,43^. The presence of enhancer-associated chromatin at this locus in fallopian-tube-derived epithelial cells is consistent with a lineage-associated regulatory element that may be retained or further activated during malignant transformation.

We further observed coordinated expression between CD47 and several neighboring genes (**Fig. S4B-F**), raising the possibility that the regulatory region participates in a broader locus-level chromatin domain rather than acting exclusively on CD47. Such coordinated expression could arise through shared enhancer activity, chromatin looping, or transcriptional coupling between neighboring genes. Importantly, these observations remain correlative, and additional experiments will be required to determine whether these genes are direct functional targets of the CD47 enhancer. Nevertheless, the observed transcriptional coordination provides a rationale for considering CD47 regulation within the context of its broader genomic neighborhood.

## Limitations of the study

This study has several limitations. Although epigenomic profiling, ChIP-qPCR, and enhancer-reporter assays support the presence of an active distal CD47 regulatory element, we did not directly perturb the endogenous enhancer, and its necessity for CD47 transcription therefore remains to be established. Similarly, while PITX2 and EP300 regulate CD47 expression and occupy this region, the precise molecular relationship between these factors, including whether PITX2 directly recruits EP300, remains unresolved. Most functional studies were performed in established ovarian cancer cell lines and macrophage co-culture models, warranting validation in patient-derived or primary HGSOC models. Finally, the presence of enhancer-associated chromatin features in fallopian tube epithelial cells is consistent with a lineage-associated regulatory program but does not establish its temporal emergence or evolutionary contribution to HGSOC development.

## Acknowledgements

We gratefully acknowledge the late Dr. Michael R. Green for his invaluable advice and generous support. We thank the UMass Chan Medical School RNAi Core Facility for providing the Brunello CRISPR library and shRNA constructs/clones; Karl Simin and the UMass Center for Clinical Sciences Biorepository, supported by NCATS (UL1-TR001453), for providing specimens; and the Deep Sequencing and Flow Cytometry Cores for providing sequencing and cell-sorting services.

## Author contributions

A.K.M, S.K.M designed research; A.K.M, S.B, T.W, A.R, A.S, AK performed research; A.K.M, T.W., S.K.M analyzed data; A.K.M, A.W, A.R, R.L and L.J.H performed bioinformatic analysis; A.K.M; S.K.M; B.A, supervised the study; and A.K.M., and S.K.M wrote the paper with the input from all authors.

## RESOURCE AVAILABILITY

### Lead Contact

Further information and requests for resources and reagents should be directed to and will be fulfilled by the Lead Contact: AKM

### Materials Availability

Plasmids and stable cell lines generated in this study are available from the Lead Contact upon reasonable request.

### Data and Code Availability

Processed data supporting the findings of this study are available within the paper and Supplementary Information. Public datasets analyzed are listed in the Key Resources Table.

## EXPERIMENTAL MODEL AND SUBJECT DETAILS

### Cell lines

OVCAR3 and SKOV3 cells were obtained from the American Type Culture Collection (ATCC), OVCAR8 cells were obtained from Creative Biolabs, PEO1 and A2780 cells were gifts from Dr. Sharon Cantor, and HEK293T cells were obtained from ATCC. Cells were maintained under sterile conditions at 37°C in a humidified incubator with 5% CO₂. OVCAR3 cells were cultured in RPMI-1640 medium, whereas OVCAR8, SKOV3, PEO1, A2780, and HEK293T cells were cultured in Dulbecco’s Modified Eagle Medium (DMEM). All culture media were supplemented with 10% heat-inactivated fetal bovine serum (FBS; Gibco), 1× non-essential amino acids (NEAA; Gibco), 1 mM sodium pyruvate (Gibco), and 100 U/mL penicillin and 100 μg/mL streptomycin (Gibco). Cells were routinely passaged at approximately 70-80% confluence using 0.25% trypsin-EDTA. experimentation.

### Human PBMCs

Peripheral blood mononuclear cells (PBMCs) were isolated from de-identified Leukopaks obtained from healthy adult donors through the Rhode Island Blood Center. Human samples were obtained and handled in accordance with applicable institutional guidelines and ethical requirements. PBMCs were isolated by Ficoll density-gradient centrifugation, and CD14⁺ monocytes were purified by magnetic bead selection. Purified monocytes were differentiated into macrophages by culture in the presence of recombinant human macrophage colony-stimulating factor (M-CSF) for 7 days.

### Cell culture conditions

All experiments were performed using exponentially growing cells maintained under sterile conditions. Stable Cas9-expressing, shRNA knockdown, and enhancer reporter cell lines were generated by lentiviral transduction followed by antibiotic selection as described in the Method Details. Unless otherwise indicated, functional assays were performed using cultures with >95% viability, and all experiments were independently repeated using separate biological replicates.

## METHOD DETAILS

### Single-cell RNA sequencing and spatial transcriptomic data analyses

Publicly available single-cell RNA sequencing (scRNA-seq) datasets from patients with high-grade serous ovarian cancer (HGSC) were analyzed using the Tumor Immune Single-cell Hub 2 (TISCH2) database, which provides uniformly processed, batch-corrected, and manually annotated tumor single-cell transcriptomic datasets^28^. The HGSC cohorts analyzed included OV_GSE118828^23^, OV_GSE130000^44^, OV_GSE147082^45^, OV_GSE151214^22^, OV_GSE154600^20^, OV_GSE158722^21^, and OV_EMTAB8107. Gene expression of CD47 was examined across malignant epithelial cells and tumor microenvironment cell populations using the standardized cell-type annotations provided by TISCH2. UMAP visualizations and normalized gene expression plots were generated through the TISCH2 web interface ( https://tisch.compbio.cn/) using the default analysis parameters. Spatial transcriptomic data from high-grade serous ovarian cancer were obtained from the Broad Institute Single Cell Portal (HGSC Spatial Cohort Discovery dataset, SCP2640, https://singlecell.broadinstitute.org/single_cell/study/SCP2640/) generated by Yeh *et al.*^46^ Spatial expression of CD47 was visualized using the interactive portal, and processed transcriptomic data were interrogated to assess the distribution of CD47 across malignant tumor regions and the surrounding tumor microenvironment.

### Epigenomic analysis

Publicly available epigenomic datasets were retrieved from the NCBI Gene Expression Omnibus (GEO). H3K27ac ChIP-seq datasets were obtained for the ovarian cancer cell lines OVCAR8, OVCA429, SKOV3, HEYA8, PEO1, and A2780. Chromatin accessibility was assessed using publicly available ATAC-seq data from OVCAR8 cells. Additional reference epigenomic datasets were included from the MCF-7 breast cancer cell line, including H3K27ac ChIP-seq, p300 ChIP-seq and BRD4 ChIP-seq datasets. Signal tracks were visualized using the Integrative Genomics Viewer (IGV) with the hg19 genome assembly across an approximately 100-kb region encompassing the CD47 locus (chr3:107,680,000–107,800,000). Regions showing enrichment of H3K27ac and increased chromatin accessibility based on ATAC-seq were used to identify active promoter and putative enhancer elements, with particular emphasis on the approximately 100-kb regulatory region surrounding the CD47 transcription start site (TSS). The NCBI GEO accession numbers for all datasets used in this analysis are provided in the Key Resources Table.

### BRD4 ChIP-seq and RNA-seq signal quantification at the CD47 locus

BRD4 ChIP-seq (DMSO, JQ1) and input control were aligned to the human genome (hg19, Bowtie, default parameters) and converted to CPM-normalized bigwig tracks. BRD4 enrichment at the CD47 proximal promoter and at the CD47 super-enhancer was calculated as log2[(mean ChIP signal + 0.01) / (mean input signal + 0.01)] over each interval, normalizing treatment signal to the matched input control rather than reporting raw coverage. CD47 gene expression was independently quantified as TPM by RSEM. As in the original study (GSE77568), RNA-seq and ChIP-seq were performed without biological replicates; consequently, bar plots report single-sample point estimates and no replicate-based statistical test was applied.

### DepMap analysis

DepMap analysis Data derived from RNA-seq (TPM) profiles across a panel of human cancer cell lines were downloaded via DepMap Public 23Q2 release (https://depmap.org/portal).

### Genome-wide CRISPR-Cas9 Knockout Screen

Cas9-expressing OVCAR8 cells were generated by lentiviral transduction with pLenti-Cas9-Blast (Addgene #52962). Lentiviral particles were produced in HEK293T cells using standard packaging plasmids, and OVCAR8 cells were transduced in the presence of 8 μg/mL polybrene. Transduced cells were selected with 5 μg/mL blasticidin for 10 days. Stable Cas9 expression was confirmed by Western blotting, and clones with high Cas9 activity were identified using a GFP-disruption reporter assay. Cas9-expressing OVCAR8 cells were subsequently transduced with a pooled lentiviral human Brunello sgRNA library targeting approximately 19,000 protein-coding genes at a multiplicity of infection (MOI) of 0.5. Following selection with puromycin (1 μg/mL) for 7 days, cells were maintained for an additional 7 days to allow CRISPR-Cas9-mediated gene editing. Cells were stained with anti-CD47-AF647 antibody (clone CC2C6), and the 5% CD47-low population was isolated by fluorescence-activated cell sorting (FACS) using a BD FACSAria Fusion cell sorter. Genomic DNA was extracted from the sorted CD47-low and unsorted populations using a Qiagen Blood Kit. sgRNA sequences were PCR-amplified and subjected to Illumina NextSeq sequencing. Sequencing data were analyzed using the CRISPR-screening pipeline established in our previous study ^47^. Briefly, raw sequencing reads were quality-controlled and mapped to the human Brunello sgRNA library, and sgRNA abundance was compared between the CD47-low and unsorted populations to identify genes associated with CD47 surface expression. Differential sgRNA representation was assessed using Fisher’s exact test, with P values adjusted for multiple testing using the Benjamini-Hochberg (BH) method.

### Lentiviral packaging and shRNA-mediated Knockdown

Two independent shRNAs targeting PITX2 or EP300 and a non-silencing control shRNA were packaged into lentiviral particles in HEK293T cells by co-transfecting the shRNA plasmid with the packaging plasmids psPAX2 and pMD2.G using Lipofectamine 3000, as previously described.^48^ Viral supernatants were collected at 48 and 72 h, filtered through a 0.45-µm membrane, and used to transduce ovarian cancer cells in the presence of 8 µg/mL polybrene. Following transduction, cells were selected with 2 µg/mL puromycin for 5-7 days to establish stable knockdown cell lines. Knockdown efficiency was confirmed by qRT-PCR and immunoblotting prior to downstream experiments

### Western Blotting and qPCR

Whole-cell protein lysates were prepared using RIPA buffer supplemented with protease and phosphatase inhibitor cocktails. Protein concentrations were determined using the Bradford Protein Assay Kit (Thermo Fisher Scientific). Equal amounts of protein (20-40 μg) were separated by SDS-PAGE, transferred onto PVDF membranes, blocked with 5% non-fat milk, and incubated with primary antibodies against CD47, PITX2, EP300, and α-tubulin (loading control), followed by HRP-conjugated secondary antibodies. Protein bands were visualized using enhanced chemiluminescence (ECL). For mRNA expression analysis, total RNA was extracted using TRIzol Reagent (Invitrogen) according to the manufacturer’s instructions. cDNA was synthesized using a reverse transcription kit, and quantitative real-time PCR (qRT-PCR) was performed using SYBR Green Master Mix (Applied Biosystems) on a QuantStudio Real-Time PCR System with gene-specific primers.

### Co-immunoprecipitation

PITX2-EP300 interaction was validated by co-immunoprecipitation (co-IP) as described previously ^49^. Briefly, cells were transfected with FLAG-PITX2 and HA-EP300 expression constructs and harvested 72 h later. Cells were lysed in non-denaturing lysis buffer supplemented with protease inhibitors, and clarified lysates were incubated overnight at 4°C with anti-FLAG or anti-HA antibodies. Immune complexes were captured using protein A beads, washed extensively, and analyzed by SDS-PAGE and immunoblotting using reciprocal antibodies. Corresponding input lysates and IgG immunoprecipitation controls were included.

### Motif Analysis of PITX2 Binding Sites in CD47 Regulatory Regions

To identify putative PITX2 binding sites within the *CD47* regulatory landscape, we performed a motif-scanning analysis using the transcription factor binding motif for human PITX2 (JASPAR ID: MA0645.1), obtained from the JASPAR 2022 database. Genomic coordinates corresponding to the *CD47* proximal promoter and a downstream enhancer region characterized by H3K27ac and ATAC-seq enrichment in OVCAR8 cells were used as input sequences. Motif scanning was conducted using the FIMO (Find Individual Motif Occurrences) tool from the MEME Suite default parameters (p-value threshold < 1e-4). PITX2 motif occurrences were visualized and annotated in the context of epigenomic features, including ATAC-seq and H3K27ac signals, to prioritize functional binding sites. Only those motif matches located within regions of open and active chromatin were considered putative regulatory PITX2 targets at the *CD47* locus.

### Chromatin Immunoprecipitation-quantitative real time PCR (ChIP-qPCR)

ChIP was performed on crosslinked OVCAR8 cells as described previously^50^, with minor modifications. Briefly, cells were fixed with 1% formaldehyde for 10 min and quenched with 125 mM glycine. Chromatin was isolated and sheared by sonication to generate fragments of approximately 200-500 bp. Immunoprecipitations were performed using 5 μg of antibodies against PITX2, EP300, or H3K27ac, with control IgG as a negative control. Following reversal of crosslinks and DNA purification, immunoprecipitated DNA was quantified by qPCR using primers targeting the CD47 distal regulatory region and promoter.

### Enhancer Reporter Assay

Candidate enhancer elements were PCR-amplified from OVCAR8 genomic DNA and cloned upstream of a thymidine kinase (TK) minimal promoter driving GFP in the sin-min-TK vector (a kind gift from *Joanna Wysocka* lab). Constructs were packaged into lentivirus and transduced into OVCAR8 cells. GFP expression was quantified by flow cytometry to assess enhancer activity.

### PBMC-Derived Macrophage Phagocytosis Assay

Peripheral blood mononuclear cells (PBMCs) were isolated from de-identified healthy donor leukopaks obtained from a blood bank using standard Ficoll-Paque PLUS (GE Healthcare) density-gradient centrifugation. CD14⁺ monocytes were subsequently purified from PBMCs by positive selection using anti-CD14 magnetic microbeads (Miltenyi Biotec) according to the manufacturer’s instructions. Monocytes were seeded at a density of 1 × 10⁶ cells/mL in RPMI-1640 medium supplemented with 10% fetal bovine serum (FBS) and 1% penicillin-streptomycin. Monocyte-to-macrophage differentiation was induced by supplementation with 50 ng/mL recombinant human M-CSF (PeproTech). Cells were maintained for 7 days, with half-media changes and re-addition of M-CSF every 3 days.For the phagocytosis assay, OVCAR8 cells expressing control or PITX2-or EP300 targeting constructs were labeled with freshly prepared pHrodo-red dye in PBS for 10 min at 37°C, washed twice, and resuspended in fresh culture medium. Labeled tumor cells were co-cultured with differentiated macrophages at a 1:1 effector-to-target (E) ratio, with 1 × 10⁵ cells of each population typically used per well in a 24-well plate. Co-cultures were maintained for 48 h at 37°C in a humidified incubator containing 5% CO₂. Following co-culture, wells were imaged using a fluorescence microscope. pHrodo-positive tumor cells/phagocytic events were quantified using ImageJ. The number of pHrodo-positive fluorescent events (red puncta) was used as a measure of macrophage-mediated phagocytosis, and the quantified data were plotted using GraphPad Prism.

### Immunohistochemical (IHC)

IHC staining was performed following a previously established protocol ^51^. De-identified formalin-fixed, paraffin-embedded (FFPE) ovarian tumor specimens were obtained from the UMass Center for Clinical and Translational Sciences Biorepository at UMass Memorial Hospital. Tissue sections (5 μm thick) were prepared from the FFPE specimens and processed for IHC using an anti-PITX2 antibody, according to the previously established staining procedure ^51^.

### Expression Correlation analysis

The mRNA expression data (RSEM-normalized TPM) for the TCGA HGSOC cohort were obtained via cBioPortal, and gene-level values were log2 (TPM+1) transformed. The association between PITX2 and CD47 expression was assessed using Spearman’s rank correlation. A two-sided Spearman correlation test was performed, and the correlation coefficient with its exact p-value is reported. For visualization, a linear regression line with 95% confidence interval was overlaid on the scatter plot.

## Supplementary figures

**Fig. S1.**
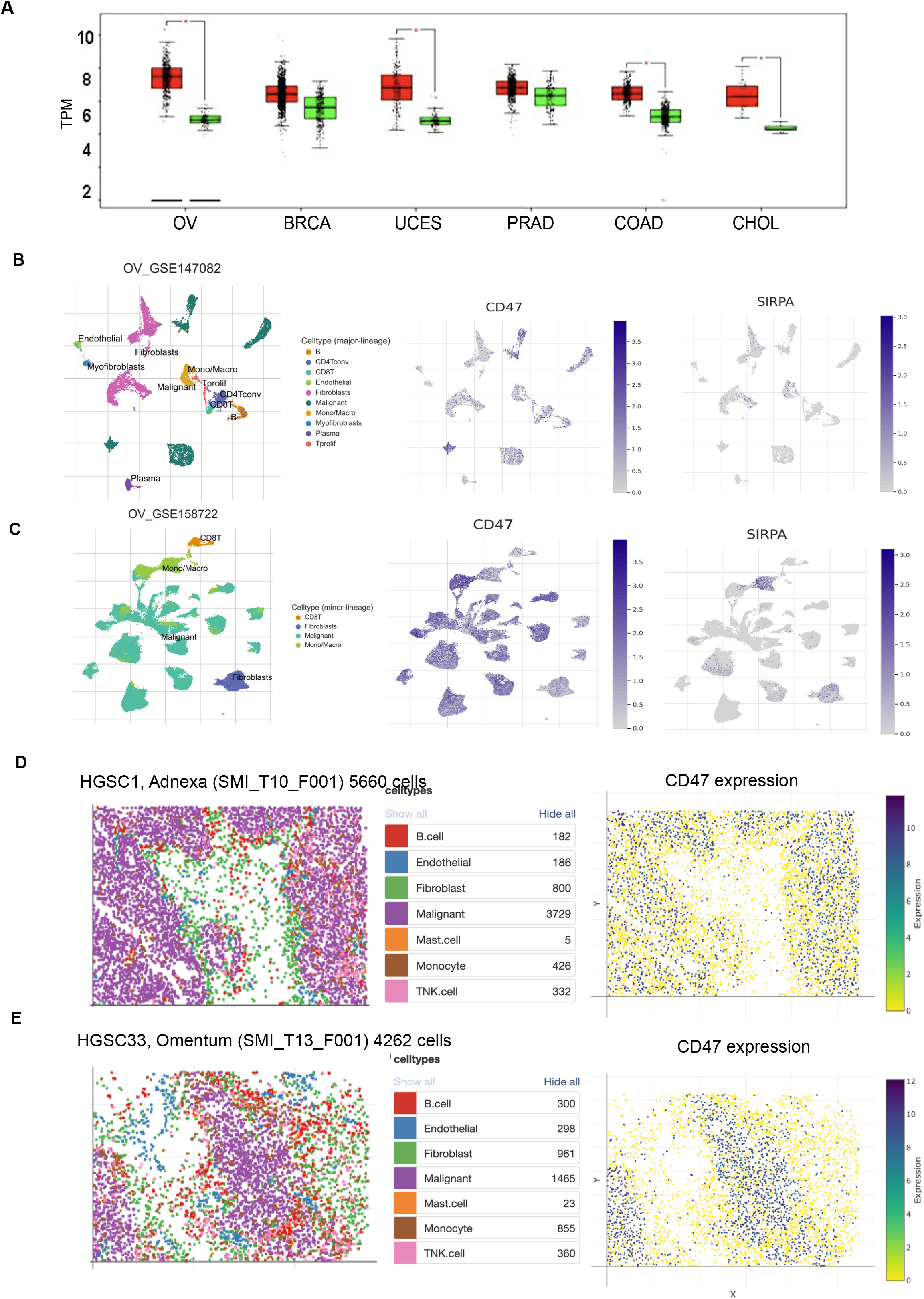
CD47 is preferentially expressed in malignant epithelial cells in HGSOC. (A) Comparison of CD47 transcript expression across TCGA cancer types, including ovarian serous cystadenocarcinoma (OV), breast invasive carcinoma (BRCA), uterine corpus endometrial carcinoma (UCEC), prostate adenocarcinoma (PRAD), colon adenocarcinoma (COAD), and cholangiocarcinoma (CHOL). Box plots show transcript abundance in TPM, with individual data points representing tumors. (B-C) UMAP visualization of scRNA-seq data from HGSOC cohorts (GSE147082 and GSE158722) colored by major cell types (left). Feature plots show expression of CD47 (middle) and its receptor SIRPA (right). CD47 is enriched in malignant epithelial cells, whereas SIRPA is predominantly expressed in monocyte/macrophage populations. (D-E) Spatial transcriptomics of two independent HGSC tumors (SMI_T10_F001 and SMI_T13_F001). Left, cell type annotation of spatial spots; right, CD47 expression. CD47 signal localizes primarily to malignant epithelial regions.

**Fig. S2.**
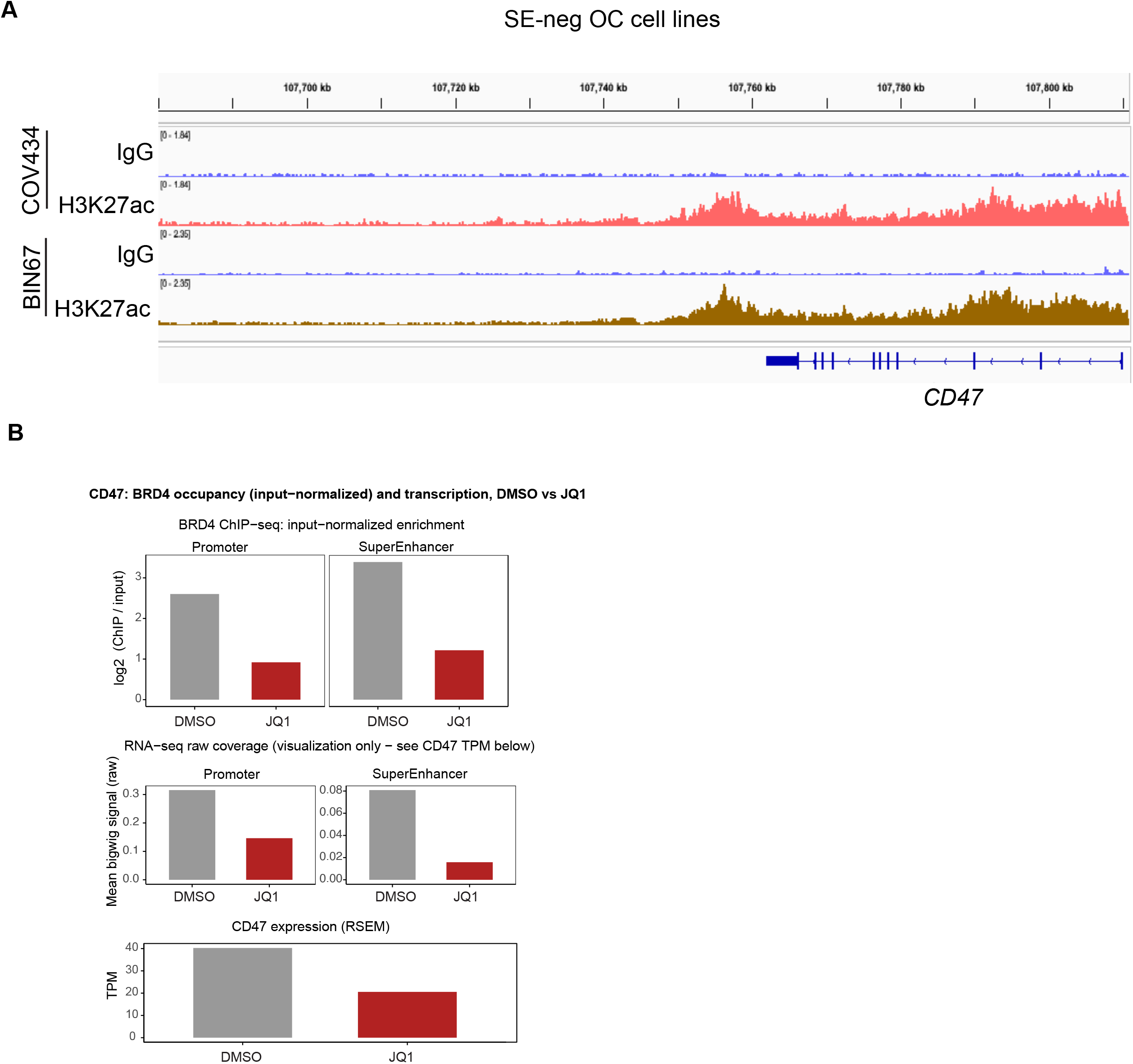
BRD4 occupancy and CD47 transcription in response to JQ1 treatment. (A) IGV tracks showing H3K27ac occupancy across the CD47 locus in BIN67 and COV434 cell lines. Promoter and super-enhancer regions are indicated. (B) BRD4 ChIP-seq occupancy at the CD47 promoter and distal super-enhancer (SE) region in DMSO-and JQ1-treated cells, shown as input-normalized enrichment. Corresponding RNA-seq coverage and CD47 transcript abundance (RSEM TPM) are shown below.

**Fig S3.**
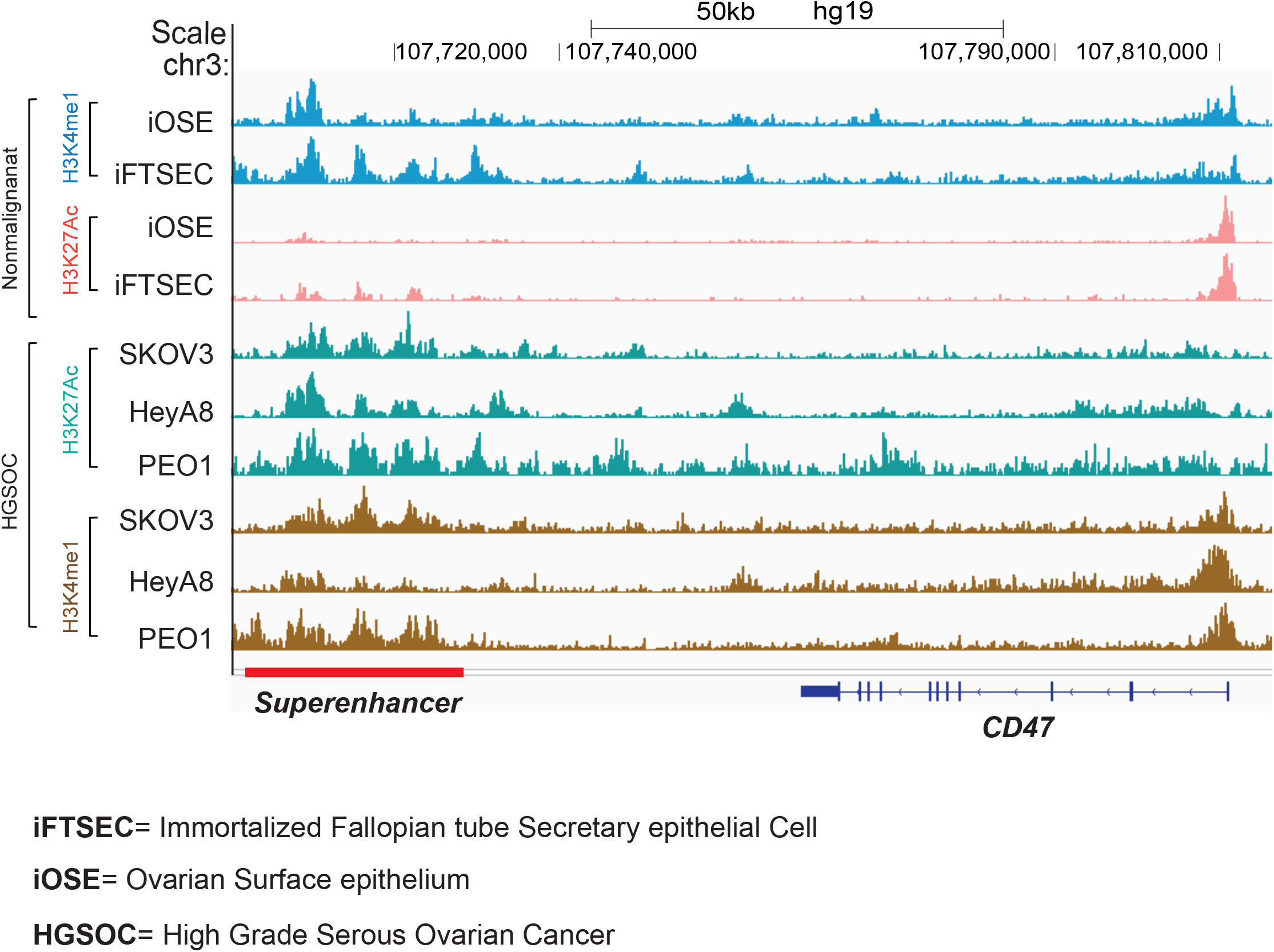
CD47-associated super-enhancer landscape across ovarian cancer and nonmalignant ovarian/fallopian tube epithelial cells. IGV tracks showing H3K4me1 and H3K27ac chromatin marks across the CD47 locus in nonmalignant immortalized ovarian surface epithelial (iOSE) and immortalized fallopian tube secretory epithelial (iFTSEC) cells, and H3K27ac and H3K4me1 signals in HGSOC cell lines SKOV3, HEYA8, and PEO1. The CD47 super-enhancer region is indicated below the genomic tracks.

**Fig S4.**
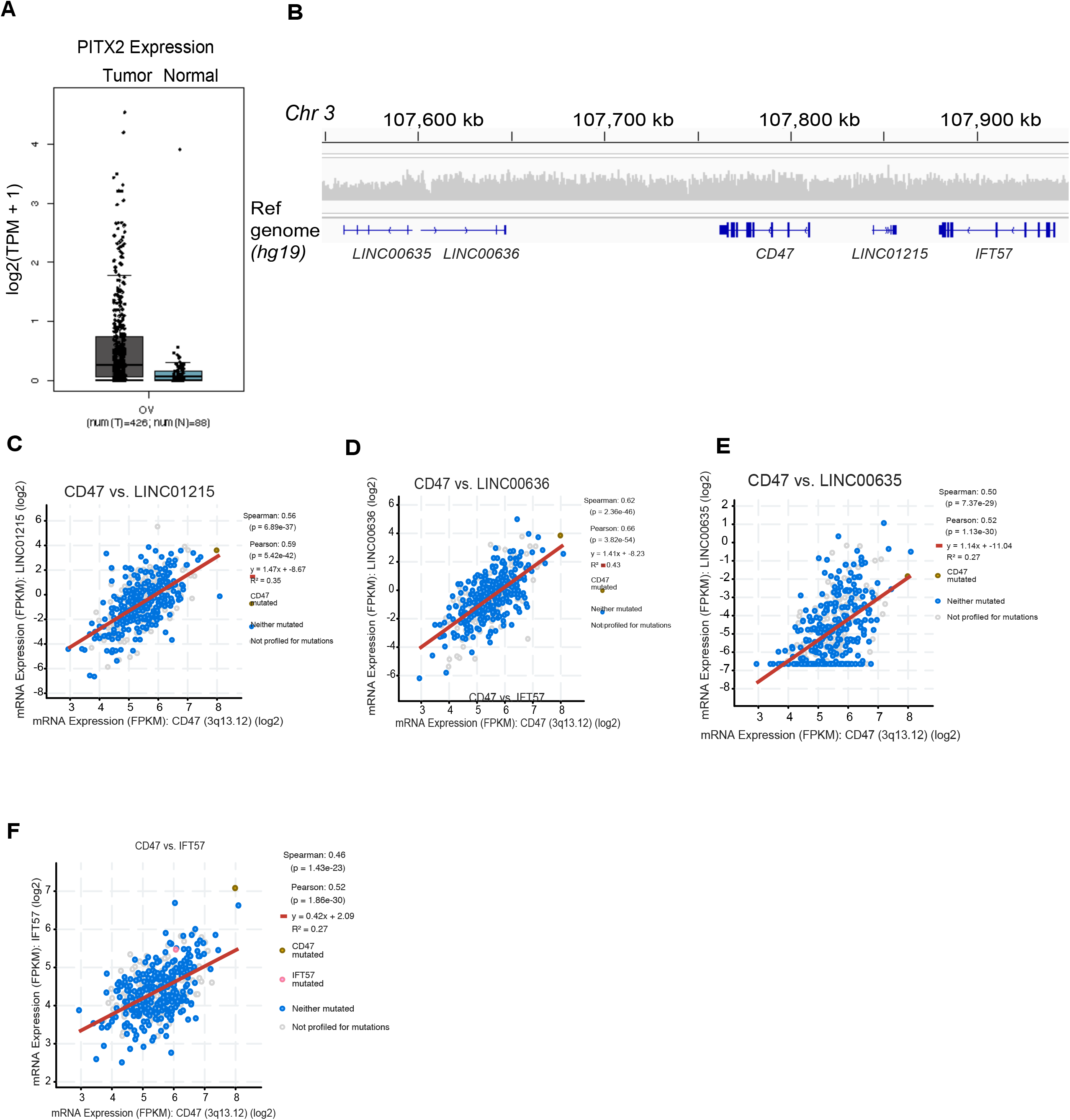
Association of CD47 expression with clinical outcome, genomic context, and neighboring gene expression in ovarian cancer. (B) Comparison of *PITX2* expression between tumor and normal ovarian tissues. (C) Genomic organization of the *CD47* locus on chromosome 3 (hg19), showing *CD47* and neighboring genes within the surrounding genomic region. (C-G) Correlation of *CD47* mRNA expression with neighboring genes *LINC01215* (C), *LINC00636* (D), *LINC00635* (E), and *IFT57* (F). Spearman and Pearson correlation coefficients, *p*-values, regression equations, and R² values are indicated in each plot.

## Supplementary data tables

**Table S1.** DepMap CD47 log2(TPM+1) Expression data

**Table S2**. Top 100 CRISPR Screening Hits

**Table S3.** Enhancer associated regulatory genes among the top 100 CRISPR screen hits.

**Table S4.** Primers and oligonucleotide used

## Key Resource Table

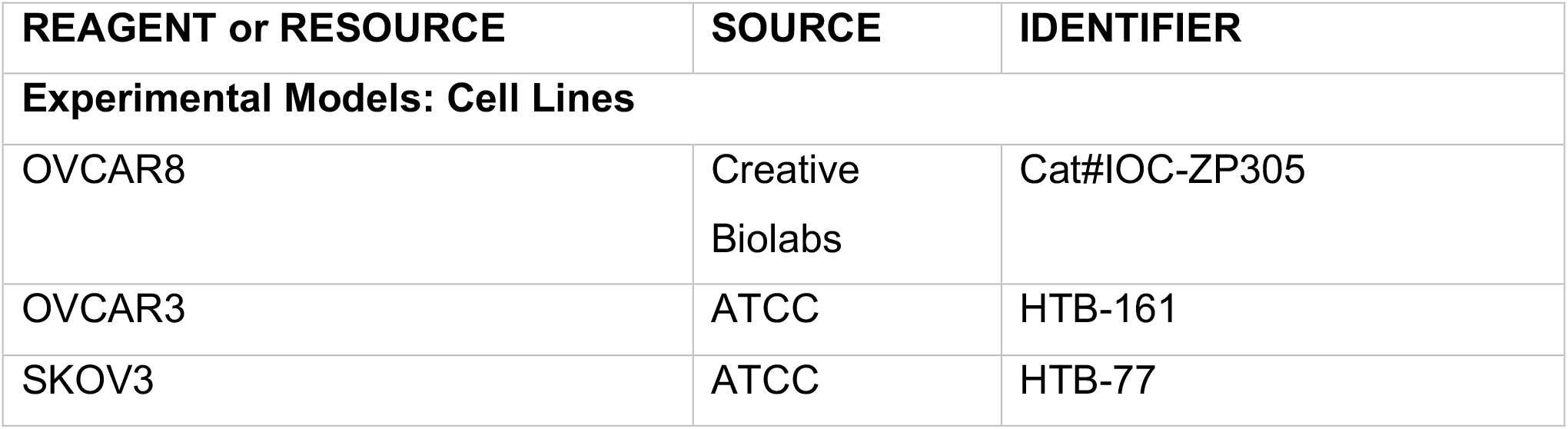
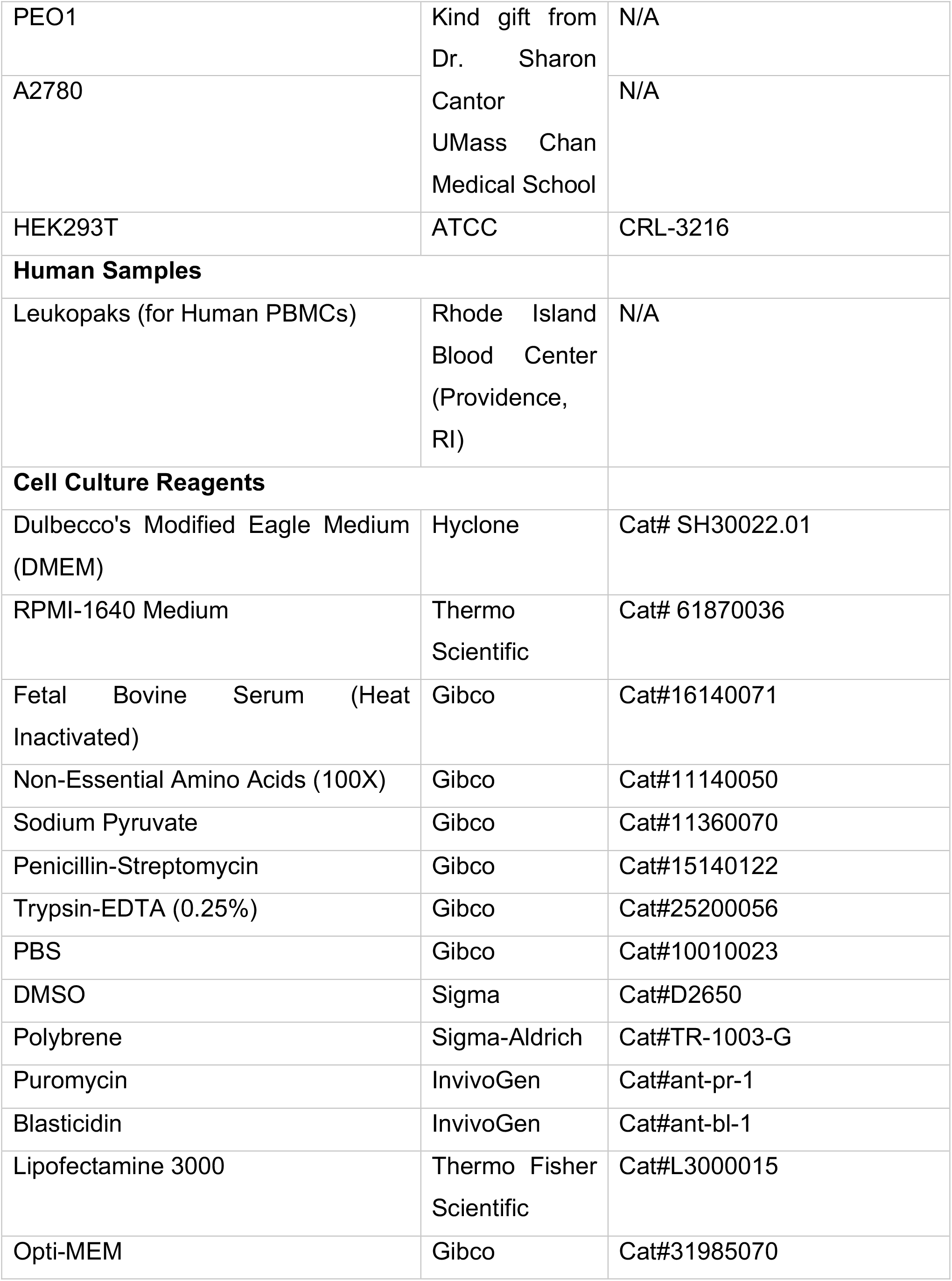
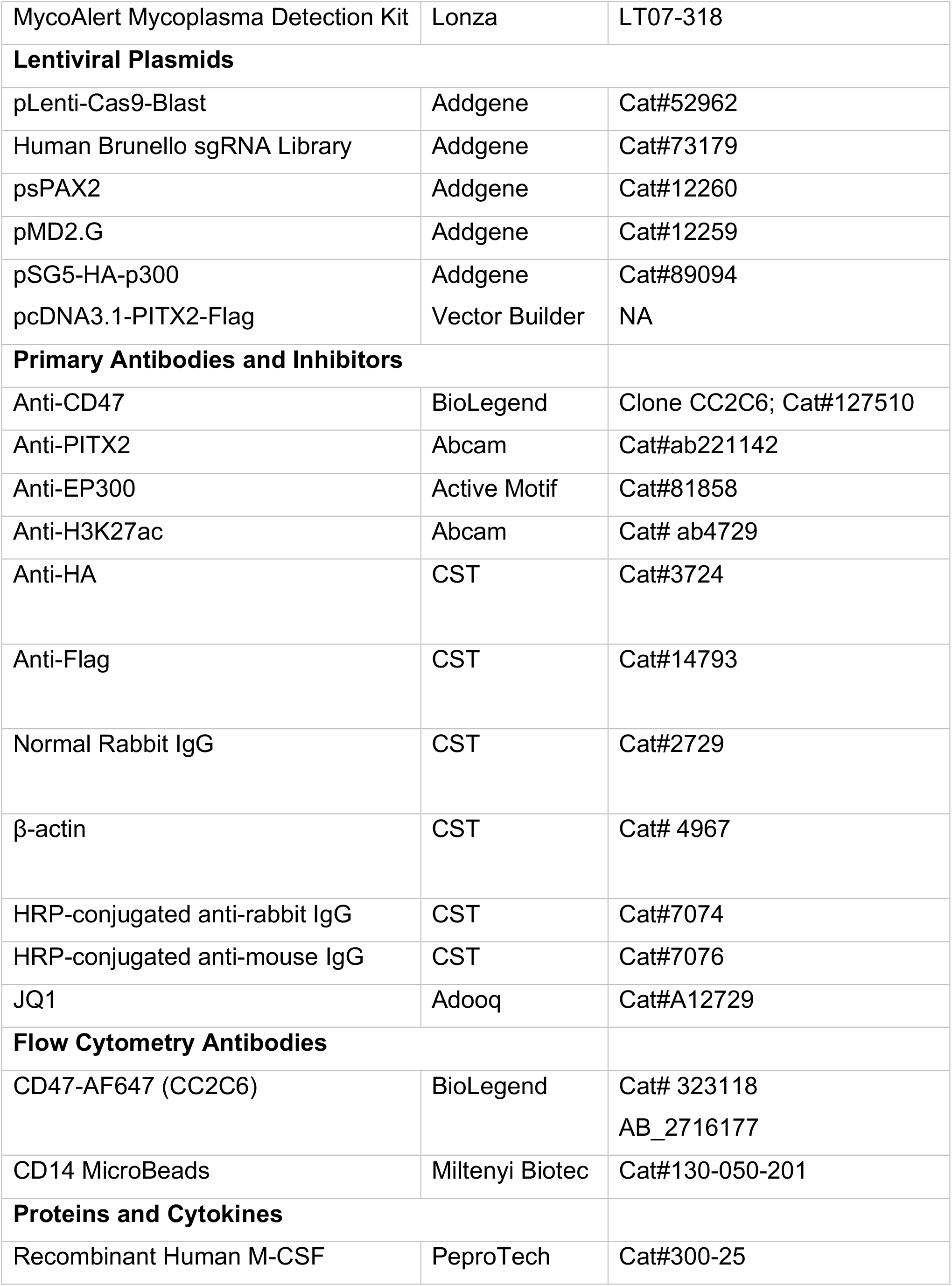
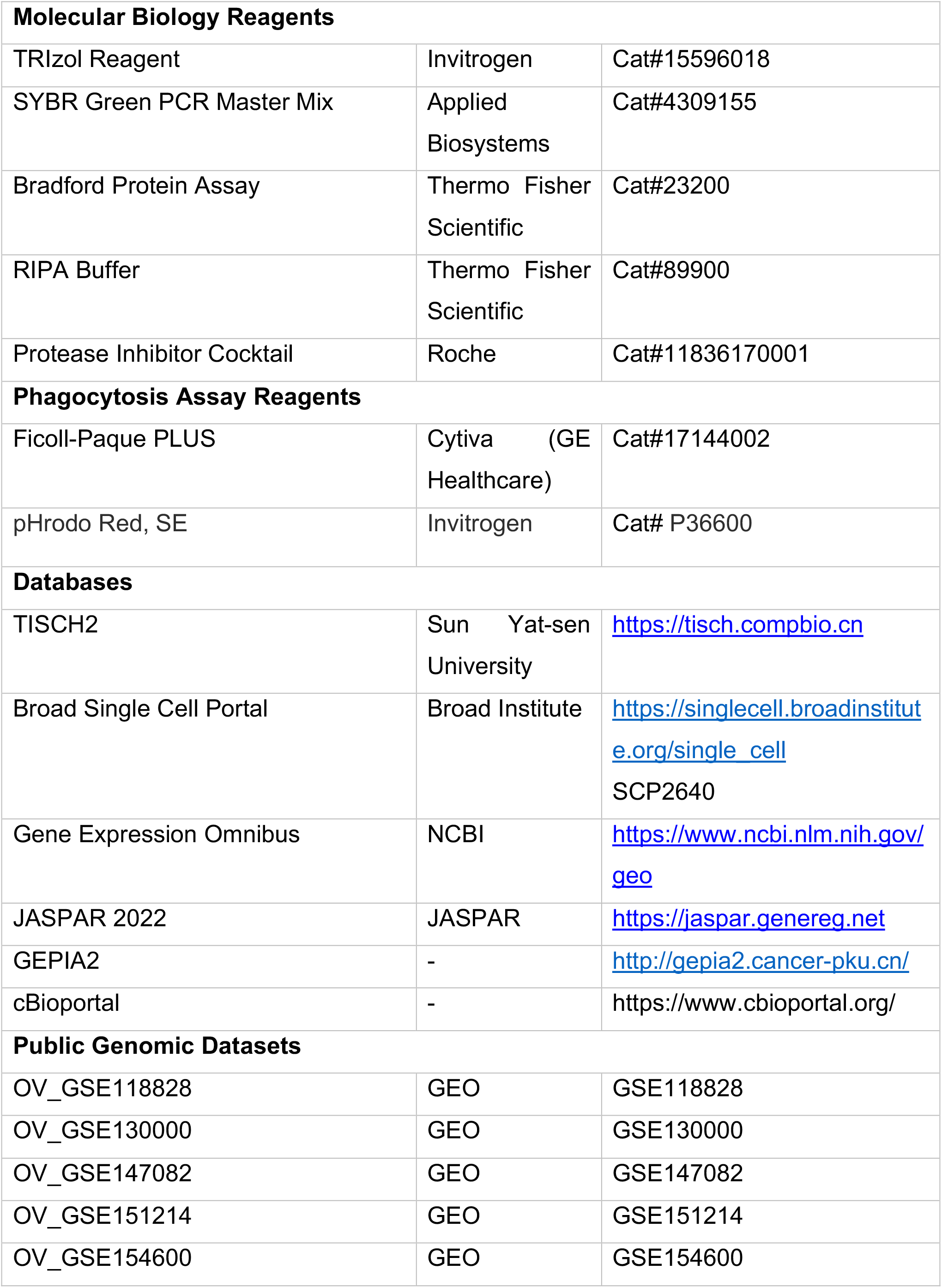
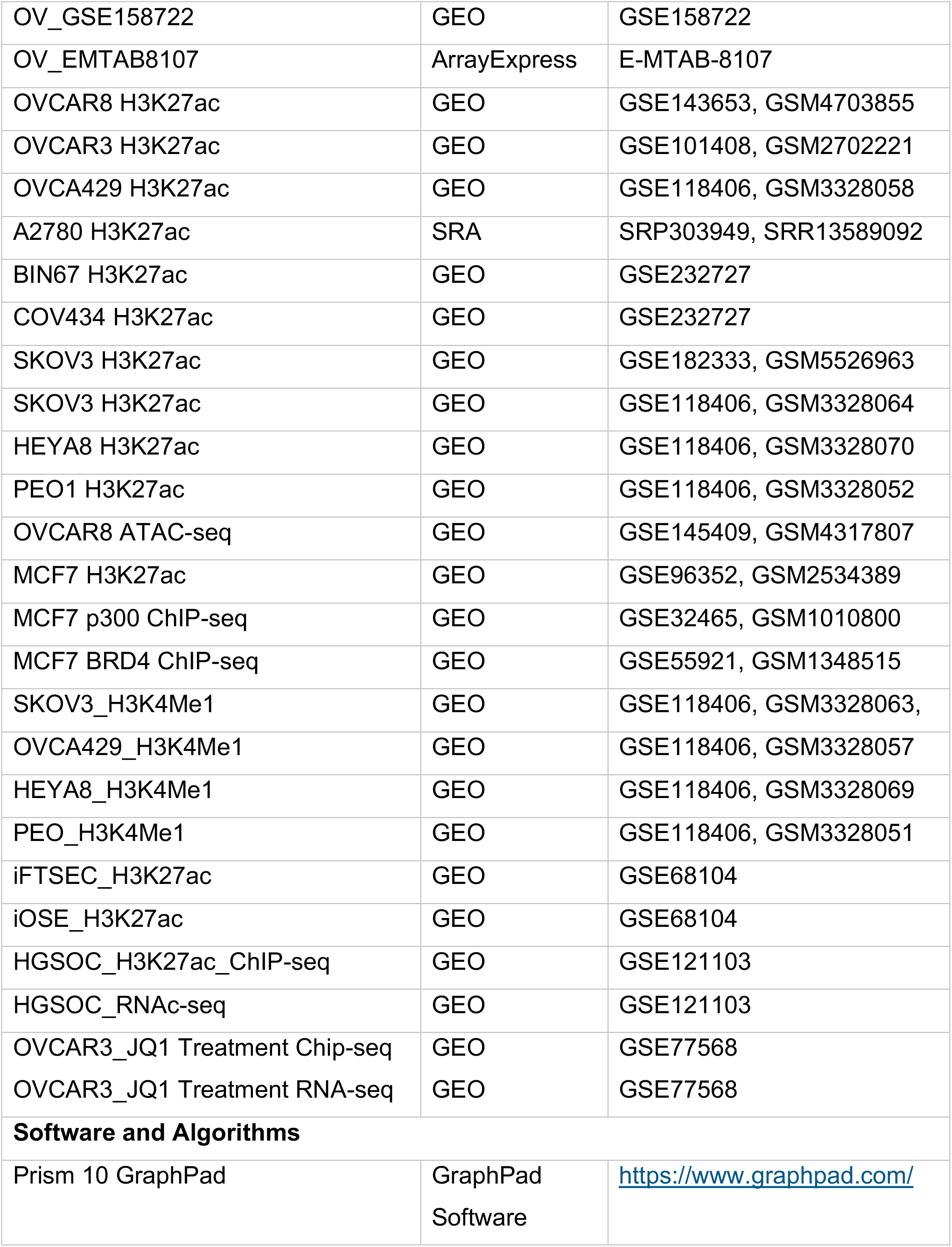
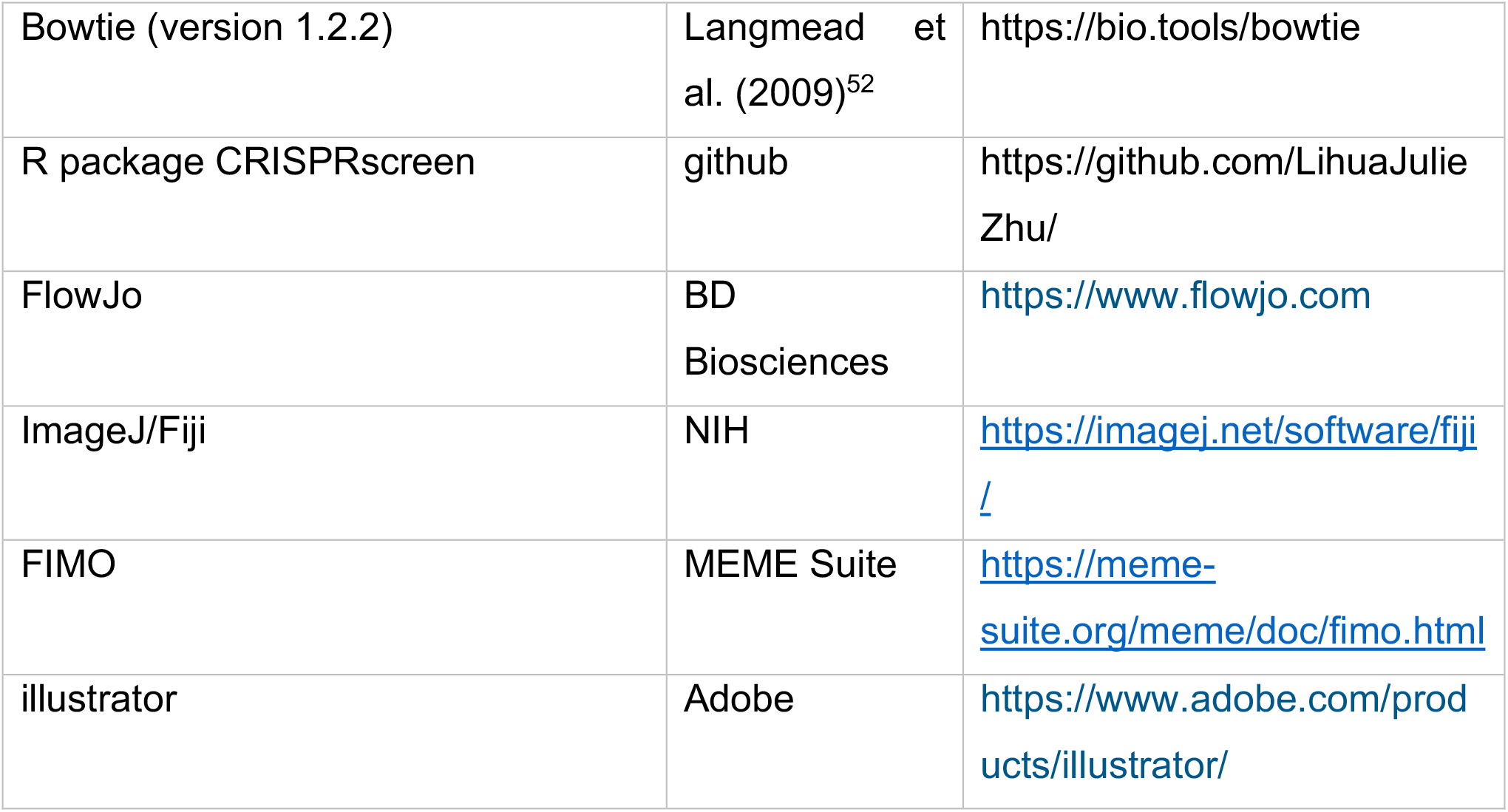

## References

1. Roerden, M., and Spranger, S. (2025). Cancer immune evasion, immunoediting and intratumour heterogeneity. Preprint at Nature Research, 10.1038/s41577-024-01111-8.

2. Galassi, C., Chan, T.A., Vitale, I., and Galluzzi, L. (2024). The hallmarks of cancer immune evasion. Preprint, 10.1016/j.ccell.2024.09.010.

3. Hanahan, D. (2022). Hallmarks of Cancer: New Dimensions. Preprint, 10.1158/2159-8290.CD-21-1059.

4. Logtenberg, M.E.W., Scheeren, F.A., and Schumacher, T.N. (2020). The CD47-SIRPα Immune Checkpoint. Preprint, 10.1016/j.immuni.2020.04.011.

5. Yang, H., Xun, Y., and You, H. (2023). The landscape overview of CD47-based immunotherapy for hematological malignancies. Preprint, 10.1186/s40364-023-00456-x.

6. Matlung, H.L., Szilagyi, K., Barclay, N.A., and van den Berg, T.K. (2017). The CD47-SIRPα signaling axis as an innate immune checkpoint in cancer. Preprint, 10.1111/imr.12527.

7. Oldenborg, P.A., Zheleznyak, A., Fang, Y.F., Lagenaur, C.F., Gresham, H.D., and Lindberg, F.P. (2000). Role of CD47 as a marker of self on red blood cells. Science (1979). 288. 10.1126/science.288.5473.2051.

8. Li, Y., Lu, S., Xu, Y., Qiu, C., Jin, C., Wang, Y., Liu, Z., and Kong, B. (2017). Overexpression of CD47 predicts poor prognosis and promotes cancer cell invasion in high-grade serous ovarian carcinoma. Am. J. Transl. Res. 9.

9. Majeti, R., Chao, M.P., Alizadeh, A.A., Pang, W.W., Jaiswal, S., Gibbs, K.D., van Rooijen, N., and Weissman, I.L. (2009). CD47 Is an Adverse Prognostic Factor and Therapeutic Antibody Target on Human Acute Myeloid Leukemia Stem Cells. Cell 138. 10.1016/j.cell.2009.05.045.

10. Liu, L., Zhang, L., Yang, L., Li, H., Li, R., Yu, J., Yang, L., Wei, F., Yan, C., Sun, Q., et al. (2017). Anti-CD47 antibody as a targeted therapeutic agent for human lung cancer and cancer stem cells. Front. Immunol. 8. 10.3389/fimmu.2017.00404.

11. Chao, M.P., Alizadeh, A.A., Tang, C., Myklebust, J.H., Varghese, B., Gill, S., Jan, M., Cha, A.C., Chan, C.K., Tan, B.T., et al. (2010). Anti-CD47 Antibody Synergizes with Rituximab to Promote Phagocytosis and Eradicate Non-Hodgkin Lymphoma. Cell 142. 10.1016/j.cell.2010.07.044.

12. Upton, R., Banuelos, A., Feng, D., Biswas, T., Kao, K., McKenna, K., Willingham, S., Ho, P.Y., Rosental, B., Tal, M.C., et al. (2021). Combining CD47 blockade with trastuzumab eliminates HER2-positive breast cancer cells and overcomes trastuzumab tolerance. Proc. Natl. Acad. Sci. U. S. A. 118. 10.1073/pnas.2026849118.

13. Chang, W.T., and Huang, A.M. (2004). α-Pal/NRF-1 Regulates the Promoter of the Human Integrin-associated Protein/CD47 Gene. Journal of Biological Chemistry 279. 10.1074/jbc.M309825200.

14. Casey, S.C., Tong, L., Li, Y., Do, R., Walz, S., Fitzgerald, K.N., Gouw, A.M., Baylot, V., Gütgemann, I., Eilers, M., et al. (2016). MYC regulates the antitumor immune response through CD47 and PD-L1. Science (1979). 352. 10.1126/science.aac9935.

15. Zhang, H., Lu, H., Xiang, L., Bullen, J.W., Zhang, C., Samanta, D., Gilkes, D.M., He, J., and Semenza, G.L. (2015). HIF-1 regulates CD47 expression in breast cancer cells to promote evasion of phagocytosis and maintenance of cancer stem cells. Proc. Natl. Acad. Sci. U. S. A. 112. 10.1073/pnas.1520032112.

16. Lo, J., Lau, E.Y.T., Ching, R.H.H., Cheng, B.Y.L., Ma, M.K.F., Ng, I.O.L., and Lee, T.K.W. (2015). Nuclear factor kappa B-mediated CD47 up-regulation promotes sorafenib resistance and its blockade synergizes the effect of sorafenib in hepatocellular carcinoma in mice. Hepatology 62. 10.1002/hep.27859.

17. Ye, Z.H., Jiang, X.M., Huang, M.Y., Xu, Y.L., Chen, Y.C., Yuan, L.W., Huang, C.Y., Yu, W.B., Chen, X., and Lu, J.J. (2021). Regulation of CD47 expression by interferon-gamma in cancer cells. Transl. Oncol. 14. 10.1016/j.tranon.2021.101162.

18. Huang, C.Y., Ye, Z.H., Huang, M.Y., and Lu, J.J. (2020). Regulation of CD47 expression in cancer cells. Preprint, 10.1016/j.tranon.2020.100862.

19. Betancur, P.A., Abraham, B.J., Yiu, Y.Y., Willingham, S.B., Khameneh, F., Zarnegar, M., Kuo, A.H., McKenna, K., Kojima, Y., Leeper, N.J., et al. (2017). A CD47-associated super-enhancer links pro-inflammatory signalling to CD47 upregulation in breast cancer. Nature Communications 8. 10.1038/ncomms14802.

20. Geistlinger, L., Oh, S., Ramos, M., Schiffer, L., LaRue, R.S., Henzler, C.M., Munro, S.A., Daughters, C., Nelson, A.C., Winterhoff, B.J., et al. (2021). Multiomic analysis of subtype evolution and heterogeneity in high-grade serous ovarian carcinoma. Cancer Res. 80. 10.1158/0008-5472.CAN-20-0521.

21. Nath, A., Cosgrove, P.A., Mirsafian, H., Christie, E.L., Pflieger, L., Copeland, B., Majumdar, S., Cristea, M.C., Han, E.S., Lee, S.J., et al. (2021). Evolution of core archetypal phenotypes in progressive high grade serous ovarian cancer. Nat. Commun. 12. 10.1038/s41467-021-23171-3.

22. Dinh, H.Q., Lin, X., Abbasi, F., Nameki, R., Haro, M., Olingy, C.E., Chang, H., Hernandez, L., Gayther, S.A., Wright, K.N., et al. (2021). Single-cell transcriptomics identifies gene expression networks driving differentiation and tumorigenesis in the human fallopian tube. Cell Rep. 35. 10.1016/j.celrep.2021.108978.

23. Shih, A.J., Menzin, A., Whyte, J., Lovecchio, J., Liew, A., Khalili, H., Bhuiya, T., Gregersen, P.K., and Lee, A.T. (2018). Identification of grade and origin specific cell populations in serous epithelial ovarian cancer by single cell RNA-seq. PLoS One 13. 10.1371/journal.pone.0206785.

24. Corona, R.I., Seo, J.H., Lin, X., Hazelett, D.J., Reddy, J., Fonseca, M.A.S., Abassi, F., Lin, Y.G., Mhawech-Fauceglia, P.Y., Shah, S.P., et al. (2020). Non-coding somatic mutations converge on the PAX8 pathway in ovarian cancer. Nat. Commun. 11. 10.1038/s41467-020-15951-0.

25. Kelly, M.R., Wisniewska, K., Regner, M.J., Lewis, M.W., Perreault, A.A., Davis, E.S., Phanstiel, D.H., Parker, J.S., and Franco, H.L. (2022). A multi-omic dissection of super-enhancer driven oncogenic gene expression programs in ovarian cancer. Nat. Commun. 13. 10.1038/s41467-022-31919-8.

26. Quintela, M., James, D.W., Garcia, J., Edwards, K., Margarit, L., Das, N., Lutchman-Singh, K., Beynon, A.L., Rioja, I., Prinjha, R.K., et al. (2023). In silico enhancer mining reveals SNS-032 and EHMT2 inhibitors as therapeutic candidates in high-grade serous ovarian cancer. Br. J. Cancer 129. 10.1038/s41416-023-02274-2.

27. Nameki, R.A., Chang, H., Yu, P., Abbasi, F., Lin, X., Reddy, J., Haro, M., Fonseca, M.A., Freedman, M.L., Drapkin, R., et al. (2023). Rewiring of master transcription factor cistromes during high-grade serous ovarian cancer development. Preprint, 10.7554/eLife.86360.1.

28. Han, Y., Wang, Y., Dong, X., Sun, D., Liu, Z., Yue, J., Wang, H., Li, T., and Wang, C. (2023). TISCH2: expanded datasets and new tools for single-cell transcriptome analyses of the tumor microenvironment. Nucleic Acids Res. 51. 10.1093/nar/gkac959.

29. Lee, Y., Miron, A., Drapkin, R., Nucci, M.R., Medeiros, F., Saleemuddin, A., Garber, J., Birch, C., Mou, H., Gordon, R.W., et al. (2007). A candidate precursor to serous carcinoma that originates in the distal fallopian tube. Journal of Pathology 211. 10.1002/path.2091.

30. Logtenberg, M.E.W., Jansen, J.H.M., Raaben, M., Toebes, M., Franke, K., Brandsma, A.M., Matlung, H.L., Fauster, A., Gomez-Eerland, R., Bakker, N.A.M., et al. (2019). Glutaminyl cyclase is an enzymatic modifier of the CD47-SIRPα axis and a target for cancer immunotherapy. Nat. Med. 25. 10.1038/s41591-019-0356-z.

31. Torlopp, A., Khan, M.A.F., Oliveira, N.M.M., Lekk, I., Soto-Jiménez, L.M. ayela, Sosinsky, A., and Stern, C.D. (2014). The transcription factor Pitx2 positions the embryonic axis and regulates twinning. Elife 3. 10.7554/eLife.03743.

32. Durbin, A.D., Wang, T., Wimalasena, V.K., Zimmerman, M.W., Li, D., Dharia, N. V., Mariani, L., Shendy, N.A.M., Nance, S., Patel, A.G., et al. (2022). EP300 Selectively Controls the Enhancer Landscape of MYCN-Amplified Neuroblastoma. Cancer Discov. 12. 10.1158/2159-8290.CD-21-0385.

33. Feng, M., Jiang, W., Kim, B.Y.S., Zhang, C.C., Fu, Y.X., and Weissman, I.L. (2019). Phagocytosis checkpoints as new targets for cancer immunotherapy. Preprint, 10.1038/s41568-019-0183-z.

34. Ryan, A.K., Blumberg, B., Rodriguez-Esteban, C., Yonei-Tamura, S., Tamura, K., Tsukui, T., De La Peña, J., Sabbagh, W., Greenwald, J., Choe, S., et al. (1998). Pitx2 determines left-right asymmetry of internal organs in vertebrates. Nature 394. 10.1038/29004.

35. Hernandez-Torres, F., Rodríguez-Outeiriño, L., Franco, D., and Aranega, A.E. (2017). Pitx2 in embryonic and adult myogenesis. Preprint, 10.3389/fcell.2017.00046.

36. Welsh, I.C., Kwak, H., Chen, F.L., Werner, M., Shopland, L.S., Danko, C.G., Lis, J.T., Zhang, M., Martin, J.F., and Kurpios, N.A. (2015). Chromatin Architecture of the Pitx2 Locus Requires CTCF-and Pitx2-Dependent Asymmetry that Mirrors Embryonic Gut Laterality. Cell Rep. 13. 10.1016/j.celrep.2015.08.075.

37. Hill, M.C., Kadow, Z.A., Li, L., Tran, T.T., Wythe, J.D., and Martin, J.F. (2019). A cellular atlas of Pitx2-dependent cardiac development. Development (Cambridge) 146. 10.1242/dev.180398.

38. Fung, F.K.C., Chan, D.W., Liu, V.W.S., Leung, T.H.Y., Cheung, A.N.Y., and Ngan, H.Y.S. (2012). Increased expression of PITX2 transcription factor contributes to ovarian cancer progression. PLoS One 7. 10.1371/journal.pone.0037076.

39. Basu, M., Bhattacharya, R., Ray, U., Mukhopadhyay, S., Chatterjee, U., and Roy, S.S. (2015). Invasion of ovarian cancer cells is induced byPITX2-mediated activation of TGF-β and Activin-A. Mol. Cancer 14. 10.1186/s12943-015-0433-y.

40. Liu, Y., Huang, Y., Fan, J., and Zhu, G.Z. (2014). PITX2 associates with PTIP-containing histone H3 lysine 4 methyltransferase complex. Biochem. Biophys. Res. Commun. 444. 10.1016/j.bbrc.2014.01.143.

41. Hilton, T., Gross, M.K., and Kioussi, C. (2010). Pitx2-dependent occupancy by histone deacetylases is associated with t-box gene regulation in mammalian abdominal tissue. Journal of Biological Chemistry 285. 10.1074/jbc.M109.087429.

42. Kim, J., Coffey, D.M., Creighton, C.J., Yu, Z., Hawkins, S.M., and Matzuk, M.M. (2012). High-grade serous ovarian cancer arises from fallopian tube in a mouse model. Proc. Natl. Acad. Sci. U. S. A. 109. 10.1073/pnas.1117135109.

43. Perets, R., Wyant, G.A., Muto, K.W., Bijron, J.G., Poole, B.B., Chin, K.T., Chen, J.Y.H., Ohman, A.W., Stepule, C.D., Kwak, S., et al. (2013). Transformation of the Fallopian Tube Secretory Epithelium Leads to High-Grade Serous Ovarian Cancer in Brca;Tp53;Pten Models. Cancer Cell 24. 10.1016/j.ccr.2013.10.013.

44. Kan, T., Zhang, S., Zhou, S., Zhang, Y., Zhao, Y., Gao, Y., Zhang, T., Gao, F., Wang, X., Zhao, L., et al. (2022). Single-cell RNA-seq recognized the initiator of epithelial ovarian cancer recurrence. Oncogene 41. 10.1038/s41388-021-02139-z.

45. Olalekan, S., Xie, B., Back, R., Eckart, H., and Basu, A. (2021). Characterizing the tumor microenvironment of metastatic ovarian cancer by single-cell transcriptomics. Cell Rep. 35. 10.1016/j.celrep.2021.109165.

46. Yeh, C.Y., Aguirre, K., Laveroni, O., Kim, S., Wang, A., Liang, B., Zhang, X., Han, L.M., Valbuena, R., Bassik, M.C., et al. (2024). Mapping spatial organization and genetic cell-state regulators to target immune evasion in ovarian cancer. Nat. Immunol. 25. 10.1038/s41590-024-01943-5.

47. Mishra, A.K., Ye, T., Banday, S., Thakare, R.P., Su, C.T.T., Pham, N.N.H., Ali, A., Kulshreshtha, A., Chowdhury, S.R., Simone, T.M., et al. (2024). Targeting the GPI transamidase subunit GPAA1 abrogates the CD24 immune checkpoint in ovarian cancer. Cell Rep. 43. 10.1016/j.celrep.2024.114041.

48. Moffat, J., Grueneberg, D.A., Yang, X., Kim, S.Y., Kloepfer, A.M., Hinkle, G., Piqani, B., Eisenhaure, T.M., Luo, B., Grenier, J.K., et al. (2006). A Lentiviral RNAi Library for Human and Mouse Genes Applied to an Arrayed Viral High-Content Screen. Cell 124. 10.1016/j.cell.2006.01.040.

49. Identification of associated proteins by coimmunoprecipitation (2005). Nat. Methods *2*. 10.1038/nmeth0605-475.

50. Banday, S., Mishra, A.K., Rashid, R., Ye, T., Ali, A., Li, J., Yustein, J.T., Kelliher, M.A., Zhu, L.J., Deibler, S.K., et al. (2025). The O-glycosyltransferase C1GALT1 promotes EWSR1::FLI1 expression and is a therapeutic target for Ewing sarcoma. Nat. Commun. 16, 1267. 10.1038/s41467-025-56632-0.

51. Chowdhury, S.R., Parikh, C.N., Kaur, A.N., DeMarco, K.D., Giwa, H.K., Mishra, A.K., Murphy, K.C., Zhou, L., Ma, B., Ye, T., et al. (2025). PPT1 is a negative regulator of STING signaling in cancer cells and its inhibition reactivates immune surveillance in cold tumors. Proc. Natl. Acad. Sci. U. S. A. 122. 10.1073/pnas.2514948122.

52. Langmead, B., Trapnell, C., Pop, M., and Salzberg, S.L. (2009). Ultrafast and memory-efficient alignment of short DNA sequences to the human genome. Genome Biol. 10. 10.1186/gb-2009-10-3-r25.

