## Supplementary material for "A Distal Super-Enhancer Regulates CD47 Expression and Promotes Macrophage-Mediated Phagocytosis of Ovarian Cancer Cells": Supplementary Table S3.docx

**Supplementary Table S3**. **Enhancer- and super-enhancer-associated regulatory genes identified among the top 100 CRISPR screen hits.**

| **Gene** | **Protein/function** | **Reported role in enhancer/super-enhancer regulation** | **Ref(s)** |
| --- | --- | --- | --- |
| **PITX2** | Homeobox transcription factor | Lineage-defining TF; recruit transcriptional activator complexes including p300 leads to hyperacetylation | [1,2] |
| **EP300** | Histone acetyltransferase | Canonical enhancer co-activator; deposits H3K27ac and activates super-enhancers | [3,4] |
| **KMT2A (MLL1)** | H3K4 methyltransferase | Establishes active enhancer chromatin (H3K4me1/3); cooperates with enhancer activation | [5] |
| **ARID2** | PBAF/SWI-SNF chromatin remodeler | Promotes enhancer accessibility and transcription factor occupancy | [6] |
| **TADA2B** | SAGA complex subunit | Component of histone acetyltransferase complex involved in enhancer activation | [7] |

**References**

**[1]**	C. Kioussi, P. Briata, S.H. Baek, D.W. Rose, N.S. Hamblet, T. Herman, K.A. Ohgi, C. Lin, A. Gleiberman, J. Wang, V. Brault, P. Ruiz-Lozano, H.D. Nguyen, R. Kemler, C.K. Glass, A. Wynshaw-Boris, M.G. Rosenfeld, Identification of a Wnt/Dvl/β-catenin → Pitx2 pathway mediating cell-type-specific proliferation during development, Cell 111 (2002). <https://doi.org/10.1016/S0092-8674(02)01084-X>.

[2]	A. Torlopp, M.A.F. Khan, N.M.M. Oliveira, I. Lekk, L.M. ayela Soto-Jiménez, A. Sosinsky, C.D. Stern, The transcription factor Pitx2 positions the embryonic axis and regulates twinning, Elife 3 (2014). <https://doi.org/10.7554/eLife.03743>.

[3]	M. Merika, A.J. Williams, G. Chen, T. Collins, D. Thanos, Recruitment of CBP/p300 by the IFNβ enhanceosome is required for synergistic activation of transcription, Mol. Cell 1 (1998). <https://doi.org/10.1016/S1097-2765(00)80028-3>.

[4]	D. Hnisz, B.J. Abraham, T.I. Lee, A. Lau, V. Saint-André, A.A. Sigova, H.A. Hoke, R.A. Young, XSuper-enhancers in the control of cell identity and disease, Cell 155 (2013). <https://doi.org/10.1016/j.cell.2013.09.053>.

[5]	K.W. Jeong, C. Andreu-Vieyra, J.S. You, P.A. Jones, M.R. Stallcup, Establishment of active chromatin structure at enhancer elements by mixed-lineage leukemia 1 to initiate estrogen-dependent gene expression, Nucleic Acids Res. 42 (2014). <https://doi.org/10.1093/nar/gkt1236>.

[6]	H.A. Malone, C.W.M. Roberts, Chromatin remodellers as therapeutic targets, Nat. Rev. Drug Discov. 23 (2024). <https://doi.org/10.1038/s41573-024-00978-5>.

[7]	S. Zhang, K. Engel, A. Fahs, C.F. Malone, K. Ross, M. Just, B. Guedes, D. Granum, K.M. Oristian, A. Kovach, G. Alexe, G. Digiovanni, L. Barbar, R. Bentley, C. Cerda-Smith, O. Le Roux, E. Mendes, S.P. Zimmerman, M. Rees, J. Roth, J.F. Shern, K.C. Wood, C.M. Counter, C.M. Linardic, K. Stegmaier, CDK8 Inhibition Releases the Muscle Differentiation Block in Fusion-driven Alveolar Rhabdomyosarcoma, (2025). <https://doi.org/10.1101/2025.07.14.663986>.
