## Supplementary material for "A Distal Super-Enhancer Regulates CD47 Expression and Promotes Macrophage-Mediated Phagocytosis of Ovarian Cancer Cells": Supplementary Table S4.docx

**Supplementary Table 4. Primers and oligonucleotide used**

| **Chip-qPCR Primers 5’-3’ Sequence** | |  |
| --- | --- | --- |
| *CD47-SE1-F*  *CD47-SE1-R* | CCTGGCTTGAAATTGGGAAAGA  TGAGCAGAGGGCCCTTGGTT | This Study |
| *CD47-SE2-F*  *CD47-SE2-R* | GGTGCATTTCATGTATGGTCAAG  GTGAAAGGGACAGGTATCACAC | This Study |
| *CD47-SE3-F*  *CD47-SE3-R1* | GGCTTTTAGACCCTTTCCCCA  CACAAAACTTTTCTTTTCTTCC | This Study |
| *CD47-TSS1-Fw*  *CD47-TSS1-Rev* | AGTCGCAGGCTCCAGACCC  AGCACGCGGACCCCAGGGGC | This Study |
| **qRT PCR Primers** | |  |
| *CD47-Fw*  *CD47-Rev*  *PITX2-Fw*  *PITX2-Rev*  *EP300-Fw*  *EP300-Rev*  *GAPDH-Fw*  *GAPDH-Rev* | CATGGCCCTCTTCTGATTTC  GGAGGTTGTATAGTCTTCTGATTGG  CGAGTCCGGGTTTGGTTCAA  GTTGGGTGGGGAAAACATGC  TCCATACCGAACCAAAGCCC  GGAGTCCACTGGAGTCTTCA  TGCACCAACTGCTTAGC  GGCATGGACTGTGGTCATGAG | [1]  [2]  [3]  [4] |
| **Enhancer Cloning Primers** | |  |
| E5-CD47_F  E5-CD47_R | TCAGACTTAGTTTGTAGATGG  ATAACACAGGGAATAGAAGC | [1] |
| **shRNA details** |  |  |
| PITX2-1  PITX2-2 | GATGCAATGATGTTTCTGAA  CCAGTCTCAACAGCCTGAAT | TRCN0000020479  TRCN0000020481 |
| EP300-1  EP300-2 | TACACTAGAGACACCTTGTAT  CCCGGTGAACTCTCCTATAAT | TRCN0000009884  TRCN0000039886 |

**References**

[1]	P.A. Betancur, B.J. Abraham, Y.Y. Yiu, S.B. Willingham, F. Khameneh, M. Zarnegar, A.H. Kuo, K. McKenna, Y. Kojima, N.J. Leeper, P. Ho, P. Gip, T. Swigut, R.I. Sherwood, M.F. Clarke, G. Somlo, R.A. Young, I.L. Weissman, A CD47-associated super-enhancer links pro-inflammatory signalling to CD47 upregulation in breast cancer, Nature Communications 8 (2017). <https://doi.org/10.1038/ncomms14802>.

[2]	H. Suzuki, X. Guo, T. Sawafuji, S. Fukuda, M. Fujiwara, K. Sasaki, T. Yamada, Y. Yamamoto, M. Kobayashi, T. Narasaka, H. Isoda, K. Tsuchiya, Transcription factor PITX2 protects intestinal epithelial cells against inflammatory stress: Implications for ulcerative colitis rectal predilection and therapeutic resistance, Biochem. Biophys. Res. Commun. 802 (2026). <https://doi.org/10.1016/j.bbrc.2026.153312>.

[3]	D.M. He, H. Wu, X.L. Wu, L. Ding, L. Xu, Y.Q. Li, The gene expression patterns of BMPR2, EP300, TGFβ2, and TNFAIP3 in B-Lymphoma cells, Cancer Biol. Med. 11 (2014). <https://doi.org/10.7497/j.issn.2095-3941.2014.03.006>.

[4]	J.X. Zhang, J.H. Wang, X.G. Sun, T.Z. Hou, GPAA1 promotes progression of childhood acute lymphoblastic leukemia through regulating c-myc, Eur. Rev. Med. Pharmacol. Sci. 24 (2020). <https://doi.org/10.26355/eurrev_202005_21182>.
